# Translation sustains productive Pol II elongation through maintenance of nuclear RNA surveillance

**DOI:** 10.64898/2026.06.21.733572

**Authors:** Mengyuan Huang, Yantao Hong, Ruiqi Deng, Yilin Liu, Yuxuan Zhou, Xiaowen Hao, Zheng Zheng, Linqing Xing, Xiaohua Shen

## Abstract

Although transcription and translation are spatially separated in eukaryotic cells, gene expression requires coordination between the nucleus and cytoplasm. Whether translation rapidly influences nuclear transcription remains unclear. Here we show that ongoing translation sustains productive RNA polymerase II (Pol II) elongation in mouse embryonic stem cells. Translation inhibition reduces nascent transcription within 15 minutes, impairs Pol II pause release, and preferentially represses long genes before substantial loss of chromatin-associated Pol II. Unexpectedly, despite reduced transcription, nuclear RNA transiently accumulates owing impaired in RNA turnover. Mechanistically, translational arrest redistributes RNA-processing and RNA-surveillance factors from the nucleus to stalled ribosome-associated complexes in the cytoplasm, reducing their nuclear availability. Acute depletion of RNA-surveillance components phenocopies elongation defects caused by translational inhibition, particularly in long genes. Together, our findings identify ongoing translation as a regulator of nuclear RNA surveillance capacity and reveal a feedback mechanism sustaining productive Pol II elongation and nuclear RNA homeostasis.

## Introduction

In eukaryotic cells, transcription and translation are spatially separated and viewed as sequential steps in gene expression. Although transcription determines the pool of RNAs available for translation, accumulating evidence suggests that translational status can also influence upstream gene-regulatory processes^1–4^. In mouse embryonic stem cells (ESCs), prolonged suppression of protein synthesis reduces active chromatin marks and suppresses nascent transcription^1^, consistent with destabilization of euchromatin regulators and transcriptional coactivators^1,5^. In contrast, translational inhibition in human cells can activate stress-response transcriptional programs and stabilize subsets of short-lived transcripts^2^, while studies in yeast have revealed rapid changes in RNA degradation that precede transcriptional adaptation^3^. Under specific physiological contexts, including nutrient stress and meiosis, translation inhibition can even promote transcription of selected gene classes^4^. However, most analyses rely on steady-state RNA measurements or long perturbations that are strongly influenced by RNA turnover and secondary cellular responses. Whether ongoing translation acutely regulates nuclear transcriptional processes remains poorly understood.

Productive transcription elongation depends not only on RNA polymerase II (Pol II) activity but also on extensive co-transcriptional RNA processing and surveillance. After promoter recruitment and pre-initiation complex assembly, RNA Pol II enters promoter-proximal pausing; gene-body elongation requires pause release by positive elongation factors such as P-TEFb/CDK9 and coordinated CTD phosphorylation, with Ser5P marking early transcription and Ser2P marking elongating Pol II^6^. During elongation, these events and nascent transcripts recruit RNA-binding proteins (RBPs), splicing factors, and RNA-surveillance machineries that promote RNA maturation, modulate premature termination, and remove aberrant RNAs^7–14^. Disruption of these pathways impairs productive elongation, particularly at transcription units that require sustained Pol II processivity and extensive processing^12,15–21^. Notably, many RNA-processing and surveillance factors shuttle between the nucleus and cytoplasm^22–26^ and interact with translating mRNPs^22,25,26^. These observations raise the possibility that ongoing translation may sustain productive transcription by maintaining the nuclear availability of RNA-surveillance machinery.

To determine whether ongoing translation rapidly influences nuclear transcriptional processes, we combined minute-scale metabolic RNA labeling, chromatin profiling, biochemical assays, and quantitative proteomics in mouse ESCs. This strategy enabled us to capture immediate consequences of translational inhibition before widespread secondary responses emerge. By integrating measurements of Pol II dynamics, RNA metabolism, and RNA regulatory factor localization, we identify an unexpected connection between translational activity, nuclear RNA surveillance, and productive transcription elongation.

## Results

### Acute translation inhibition leads to rapid genome-wide transcriptional repression

To determine whether ongoing translation acutely influences nuclear transcription, we inhibited translation in ESCs using mechanistically distinct inhibitors: cycloheximide (CHX), harringtonine (HAR), and anisomycin (ANS), which target translational elongation, initiation, and peptide-bond formation, respectively. Short Homopropargyl Glycine (HPG, an amino acid analogue) pulses confirmed effective translational inhibition, revealing marked reductions in translation-associated signals in both whole cells and isolated nuclei following CHX or HAR treatment (Fig. 1A and Supplementary Fig. 1 A-D). We next asked whether ongoing translation is acutely required for nuclear transcription.

**Figure 1.**
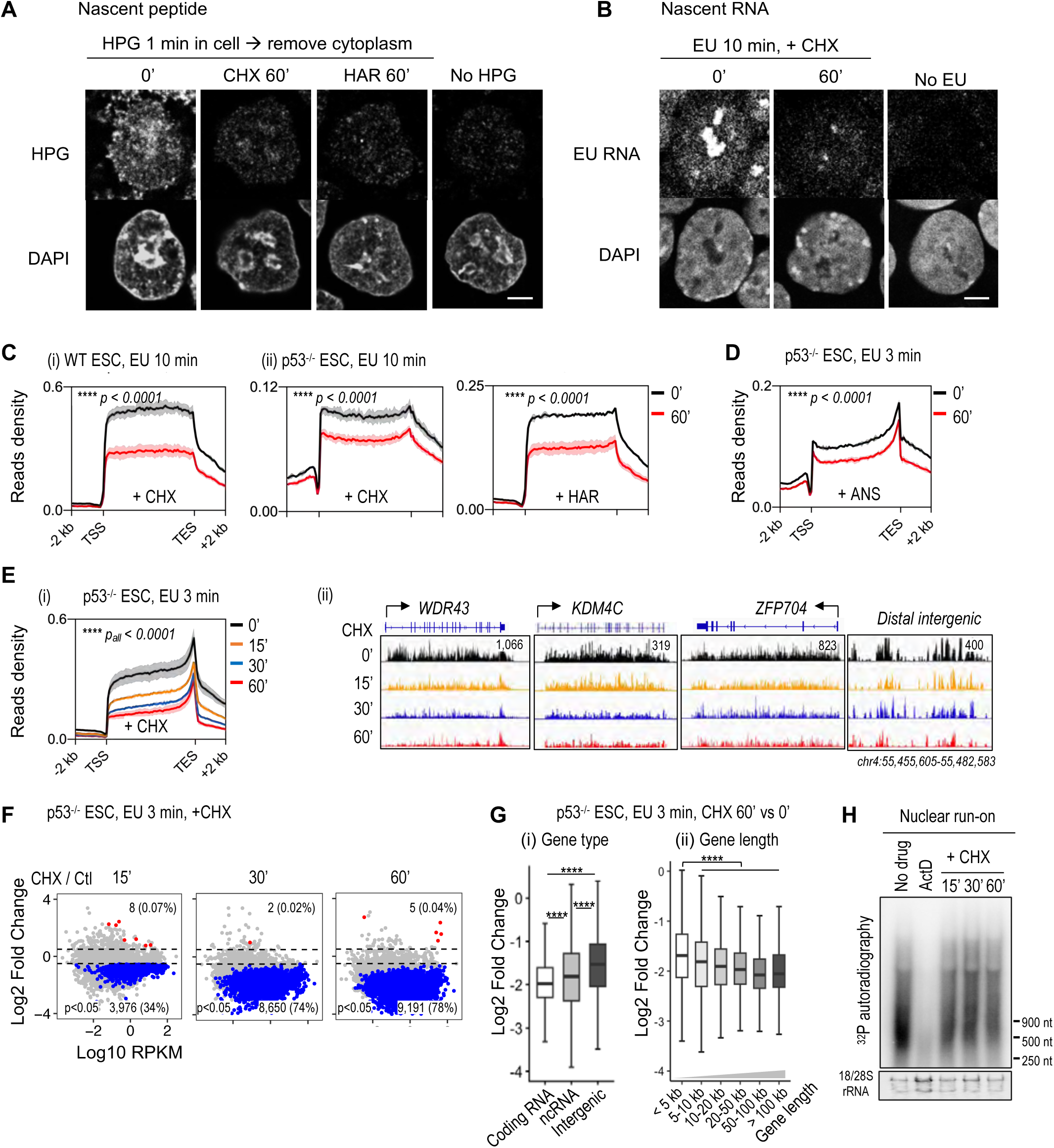
Rapid impairment of productive transcription upon translation inhibition. (A) HPG incorporation for detecting translational output. Cells were labeled with HPG for 1 min, followed by on-plate nuclear extraction, fixation, click chemistry, and fluorescence imaging. Quantification was shown in Supplementary Fig. 1C. Scale bar: 5 *μ*m. (B) EU incorporation for detecting transcriptional output. Cells were labeled with EU for 10 min on plate, fixation, click chemistry, and fluorescence imaging. Quantification is shown in Supplementary Fig. 1E. Scale bar: 5 *μ*m. (C) Metagene analysis of cell-number normalized EU-seq (10 min labeling) across 22,470 mouse protein-coding gene bodies with ±2 kb flanking regions. Wild-type and p53^-/-^ ESCs were treated with CHX (10 *μ*g/mL) or HAR (20 *μ*M) for 60 min. Unless otherwise noted, CHX and HAR were applied at identical concentrations in all subsequent experiments. Only the sense strand is shown. Solid lines indicate mean of two biological replicates and shaded areas represent SEM. Statistical significance was assessed using a two-sided Kolmogorov-Smirnov test. Spike-in controls were used for normalization. (D) Metagene profiles of cell-number normalized EU-seq (3 min labeling) across 22,470 mouse protein-coding gene bodies with ±2 kb flanking regions. p53^-/-^ ESCs were treated with translation inhibitor ANS (1 *μ*g/mL) for 60 min. Unless otherwise noted, ANS was applied at identical concentrations in all subsequent experiments. Only the sense strand is shown. Solid lines indicate mean values of two biological replicates and shaded areas represent SEM. Statistical significance was assessed using a two-sided Kolmogorov-Smirnov test. Spike-in controls were used for normalization. (E) (i) Metagene profiles of cell-number normalized EU-seq (3 min labeling) across 22,470 mouse protein-coding gene bodies with ±2 kb flanking regions following CHX treatment for 15-60 min. Solid lines indicate mean values of two biological replicates and shaded areas represent SEM. Only sense strand is shown. Statistical significance was assessed using a two-sided Kolmogorov-Smirnov test. Spike-in controls were used for normalization. Statistical comparisons shown are between CHX-treated and control groups. (ii) Representative IGV snapshots including *WDR43*, *KDM4C*, *ZFP704*, and an intergenic region of 3-min labeled EU-seq. (F) Volcano plots showing expression changes of cell-number normalized EU-seq (3 min labeling). The x axis: log₂ expression levels in control cells. The y axis: log₂ fold changes of genes with detectable expression (RPKM > 0). Genes with p < 0.05 and |log₂ fold change| > 0.5 (dashed lines) are shown as significantly upregulated (red) or downregulated (blue), with non-significant genes shown in gray. Gene significance was assessed using a two-sided t test from two biological replicates. (G) Box plots showing EU-seq signal changes (3 min labeling) before and after 60 min CHX treatment across genomic regions (i, coding RNA, ncRNA and intergenic bins, n = 10,019/467/4,272) and gene length groups (ii, <5, 5-10, 10-20, 20-50, 50-100, >100 kb, n = 2,374/1,729/2,782/4,113/2,276/2,426). Boxes indicate the 25^th^-75^th^ percentiles with medians, and whiskers denote the 5^th^ -95^th^ percentiles. Statistical significance was assessed using a two-sided Wilcoxon rank-sum test. Statistical comparisons in (ii) were made between each group and the first group (<5 kb). (H) Nuclear run-on assay using ^α-32P^UTP. *18/28S* rRNA signal are shown below. Quantification is shown in Supplementary Fig. 1I. Significance: ns (>0.05), * (≤0.05), ** (≤0.01), *** (≤0.001), **** (≤0.0001). The same criteria were applied to all analyses shown in this study.

Fluorescence imaging of 5-ethynyl uridine (EU)-labeled RNA revealed widespread reduction in nascent RNA throughout the nucleoplasm and nucleolus after 60 min of CHX treatment (Fig. 1B and Supplementary Fig. 1E). Cell-number-normalized EU-seq using a 10-min labeling window confirmed a global reduction in nascent transcription (Fig. 1C, i). Similar effects were observed with HAR or ANS treatment and were maintained in p53-deficient ESCs (Fig. 1C, ii and Fig. 1D), indicating that this response is independent of p53 activation.

Time-resolved EU-seq showed that transcriptional repression emerged rapidly. A 3-min EU pulse detected an ∼30% reduction in gene-body transcription within 15 min of CHX treatment, which became more pronounced over 30–60 min (Fig. 1E). The number of significantly downregulated genes increased from 3,976 at 15 min to 9,191 at 60 min, whereas relatively few genes showed increased transcription (Fig. 1F). Thus, ongoing translation is continuously required to sustain genome-wide transcriptional output.

Transcriptional repression was not uniform across the genome. Protein-coding genes were most strongly affected, followed by long noncoding RNAs (lncRNAs), whereas distal intergenic transcription (>10 kb from annotated genes) exhibited comparatively lower sensitivity (Fig. 1G and Supplementary Fig. 1F). Sensitivity increased with gene length, with long genes exhibiting the strongest reduction in nascent transcription (Fig. 1G and Supplementary Fig. 1G, H), suggested that productive elongation may represent an early target of translational inhibition.

To assess whether the rapid transcriptional reduction reflects changes intrinsic to the nucleus, we performed nuclear run-on assays in isolated nuclei following CHX treatment. In the presence of sarkosyl, which blocks new Pol II initiation, this assay selectively measures pre-engaged polymerase activity. Actinomycin D abolished run-on signals, whereas CHX reduced run-on activity by ∼40–60% within 15–60 min (Fig. 1H and Supplementary Fig. 1I). Because transcription was measured ex vivo, these results indicate that translation inhibition alters the nuclear environment in a manner that persists outside the cytoplasm, impairing the activity of engaged polymerases.

### Translation inhibition uncouples Pol II occupancy from productive elongation

The preferential sensitivity of long genes suggested that translational inhibition may impair productive elongation rather than transcription initiation. To directly examine RNA Pol II behavior, we quantified genome-wide Pol II occupancy and activity following CHX treatment. Unexpectedly, transcriptional output declined before substantial loss of chromatin-associated Pol II. At 15 min, gene-body Pol II occupancy remained largely unchanged (Fig. 2A) despite a marked reduction in nascent RNA synthesis. Thus, engaged polymerases persisted on chromatin even as transcriptional output decreased. By 30–60 min, Total, Ser2P, and Ser5P Pol II occupancy progressively declined across gene bodies and transcription end sites (TES), whereas promoter-proximal occupancy was preserved (Fig. 2A, B and Supplementary Fig. 2A, B).

**Figure 2.**
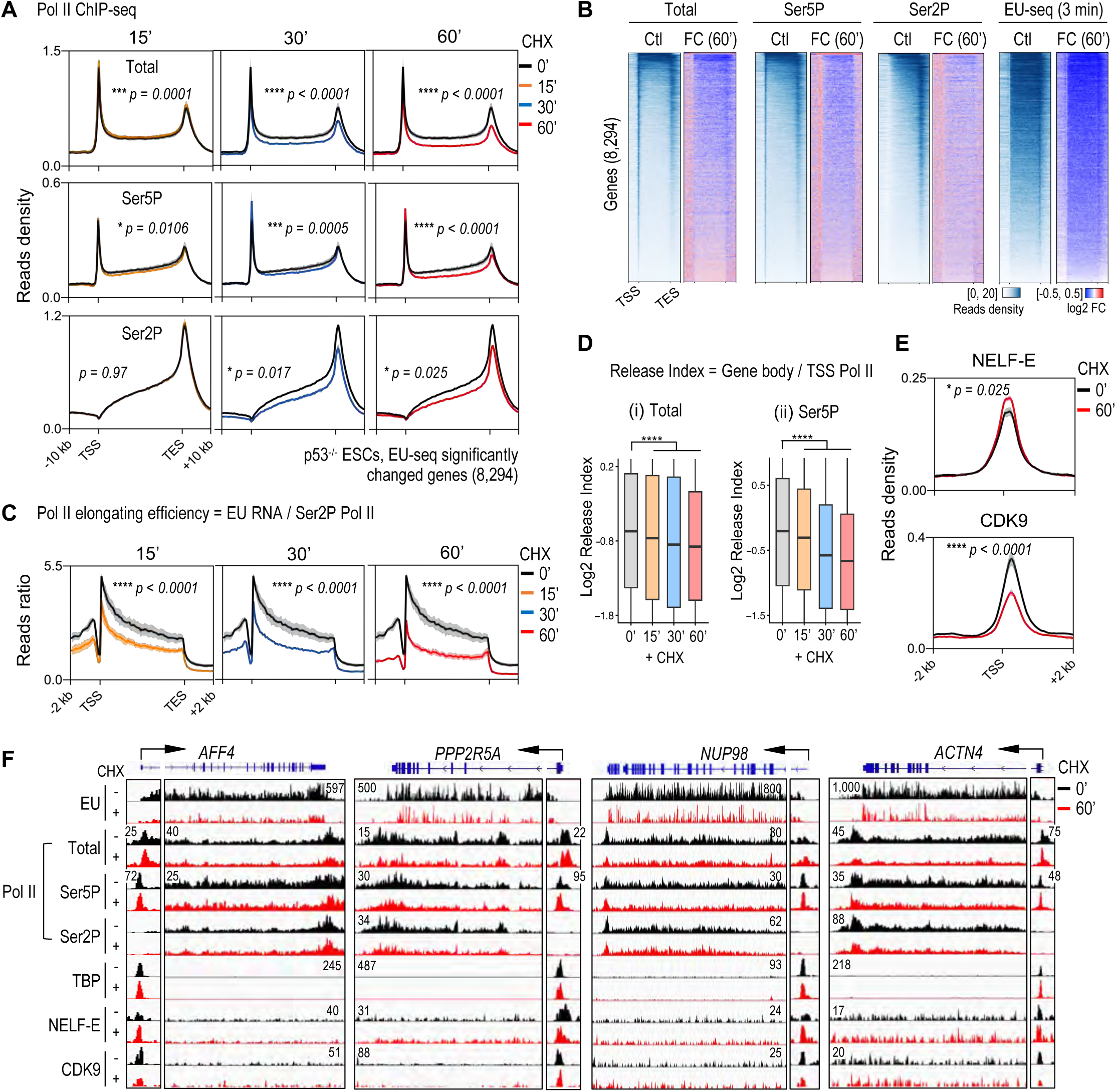
Rapid decoupling of Pol II occupancy and productive elongation upon translation inhibition. (A) Metagene analysis of Total, Ser5P and Ser2P Pol II ChIP-seq signals across 8,294 CHX-responsive genes (significantly changed during 15-60 min CHX treatment). Profiles are shown across gene bodies with ±10 kb flanking regions. Solid lines indicate mean values of two biological replicates and shaded areas represent SEM. Statistical significance was assessed using a two-sided Kolmogorov-Smirnov test. Signals were normalized by sequencing depth. (B) Heatmap showing Pol II distribution across 8,294 CHX-responsive genes and their ±2 kb flanking regions. Signals represent the mean of two biological replicates and are shown as either Z-score normalized values or log₂ fold change. Genes or bins with no detectable reads across all conditions were excluded. (C) Metagene analysis of Pol II elongation efficiency, calculated as EU-seq signal divided by Ser2P Pol II occupancy, across 8,294 CHX-responsive genes and their ±2 kb flanking regions. Solid lines indicate mean values of two biological replicates and shaded areas represent SEM. Statistical significance was assessed using a two-sided Kolmogorov-Smirnov test. Signals were normalized by sequencing depth. (D) Box plots showing release index, calculated as gene body/TSS ratio of Total and Ser5P Pol II across expressed genes (FPKM>0) (n = 7,553 and 8,140, respectively). Boxes represent the 25^th^-75^th^ percentiles with medians indicated, and outliers are omitted. Statistical significance was assessed using a two-sided Wilcoxon rank-sum test. (E) Metagene profiles of NELF-E and CDK9 ChIP-seq signals across 8,294 CHX-responsive genes and their ±2 kb flanking regions. Solid lines indicate mean values of two biological replicates and shaded areas represent SEM. Statistical significance was assessed using two-way ANOVA for NELF-E or a two-sided Kolmogorov-Smirnov test for CDK9. Signals were normalized by sequencing depth. (F) Representative IGV snapshots showing EU-seq (3 min labeling) and Total, Ser5P, and Ser2P Pol II signals at *AFF4*, *PPP2R5A*, *NUP98*, and *ACTN4* loci.

To quantify the functional consequence, we defined elongation efficiency as the ratio of nascent RNA production to chromatin-associated Ser2P Pol II. Elongation efficiency was significantly reduced starting at 15 min of CHX treatment (Fig. 2C), indicating that translational inhibition compromises the activity of engaged polymerases before detectable polymerase loss. Consistent with this, the release index of Total and Ser5P Pol II, defined as the ratio of Pol II density across gene bodies relative to TSS regions, progressively decreased over time, reflecting impaired transition into productive elongation (Fig. 2D). Concordantly, the negative elongation factor NELF-E accumulated at promoters, whereas the positive elongation kinase CDK9 declined from chromatin (Fig. 2E), shifting the balance toward a paused transcriptional state.

Total protein levels of Pol II, TBP, NELF-E, and CDK9 were stable (Supplementary Fig. 2C), and TBP occupancy at promoters remained largely unchanged (Supplementary Fig. 2D-E), consistent with the notion that translational inhibition primarily affects elongation rather than initiation (Fig. 2F). Together, these findings reveal an early uncoupling of Pol II occupancy from transcriptional output, indicating that engaged polymerases lose functional competence before dissociating from chromatin.

### Nuclear RNA accumulates through coordinated suppression of RNA synthesis and decay

Despite rapid declines in gene-body transcription using short 3-min EU pulses, longer 10-min EU-labeling windows revealed an unexpected transient increase in nuclear RNA following CHX treatment (Fig. 3A, B). This discrepancy suggested that translational inhibition affects not only RNA synthesis but also nuclear RNA turnover.

**Figure 3.**
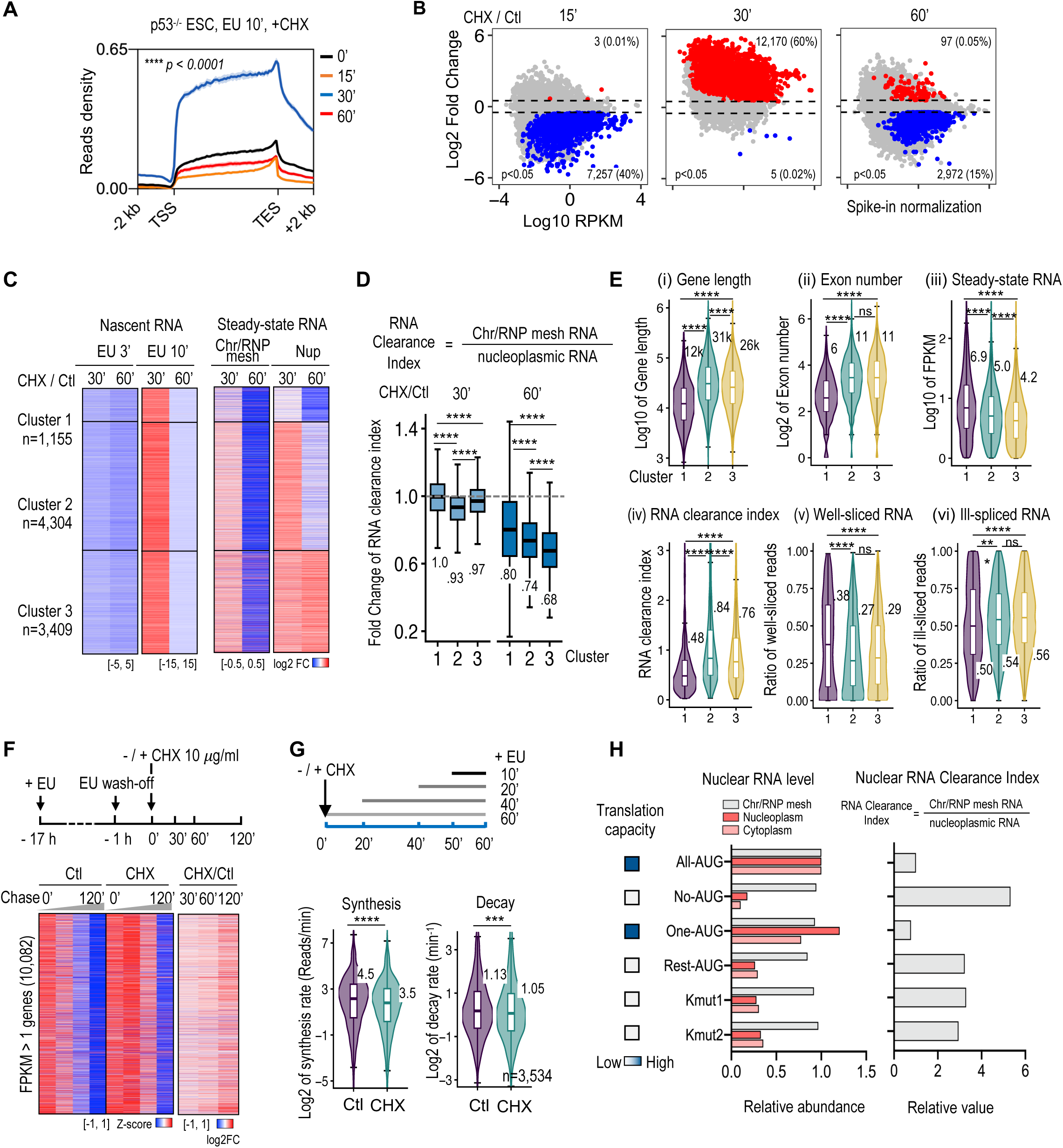
Nuclear RNA transient accumulation associated with suppression of RNA synthesis and turnover. (A) Metagene profiles of cell-number normalized EU-seq (10 min labeling) across 22,470 mouse protein-coding gene bodies with ±2 kb flanking regions. Solid lines indicate mean values of two biological replicates and shaded areas represent SEM. Only sense strand is shown. Statistical significance was assessed using a two-sided Kolmogorov-Smirnov test. Signals were normalized by cell number. Statistical comparisons shown are between CHX-treated and control groups. (B) Volcano plots showing expression changes of cell-number normalized EU-seq (10 min labeling). The x axis: log₂ expression levels in control cells. The y axis: log₂ fold changes of genes with detectable expression (RPKM > 0). Genes with p < 0.05 and |log₂ fold change| > 0.5 (dashed lines) are shown as significantly upregulated (red) or downregulated (blue), with non-significant genes shown in gray. Statistical significance was assessed using a two-sided t test from two biological replicates. (C) Heatmap showing absolute log₂ fold changes between CHX treatment and control. Left panel shows nascent RNA changes measured by EU-seq (3- or 10-min labeling). Right panel shows nuclear steady-state RNA changes in chromatin/RNP mesh (Chr/RNP mesh) and nucleoplasmic (Nup) fractions. Genes with FPKM > 1 (n = 8,868) were included. Genic (EU 10 min) or intronic (EU 3 min) reads were used as a proxy for nascent transcription, whereas exonic reads were used to quantify steady-state RNA abundance. Genes were grouped into three clusters based on their nuclear RNA change patterns. Cluster 1 contained 1,155 genes and showed limited nucleoplasmic RNA accumulation. In contrast, clusters 2 and 3 represented two major response classes, comprising 4,304 and 3,409 genes, respectively, that accumulated nucleoplasmic RNA despite reduced nascent transcription. This retention was transient in cluster 2 but persisted through 60 min in cluster 3. (D) Boxplot showing nuclear RNA clearance index, defined as the ratio of Chr/RNP mesh to nucleoplasmic (Nup) RNA levels. Clusters were defined as in Fig. 3C. Boxes indicate the 25^th^ -75^th^ percentiles with medians shown, and whiskers represent the 5^th^ -95^th^ percentiles. Outliers were omitted. Numbers below boxes indicate median values for each group. Statistical significance was assessed using a two-sided Wilcoxon rank-sum test. (E) Violin and box plots showing distributions of gene length, exon number, steady-state RNA abundance, RNA clearance index, and splicing ratio across three gene groups defined in Fig. 3C. Boxes indicate the 25^th^ -75^th^ percentiles with medians shown, and whiskers represent the 5^th^ -95^th^ percentiles. Outliers were omitted. Statistical significance was assessed using a two-sided Wilcoxon rank-sum test. (F) Schematic and heatmap showing RNA stability measured by metabolic EU pulse-chase experiments across 10,082 genes with FPKM > 1. Exonic read counts per gene were normalized to rRNA, normalized to the 0 min time point, and Z-score transformed for visualization. The right panel shows log₂ fold changes in read counts at each chase time point relative to 0 min chase time. (G) Nuclear RNA synthesis and degradation rates inferred using INSPEcT. Upper: Schematic of EU metabolic labeling (10 min to 60 min) with or without CHX treatment (60 min). Lower: Violin plots show distributions of 3,534 genes with detectable 10-min EU signal (RPKM > 1). Boxes indicate the 25^th^ -75^th^ percentiles with medians shown, and whiskers represent the 5^th^ -95^th^ percentiles. Outliers were omitted. Statistical significance was assessed using a two-sided Wilcoxon rank-sum test. (H) â-globin reporter constructs with altered translation capacity were generated by mutating the open reading frame, including disruption of AUG start codons and the Kozak sequence (see also Supplementary Fig. 3F). The density of blue squares reflects translational capacity. Constructs were stably transfected into ESCs. Following subcellular fractionation, reporter RNA levels in chromatin/RNP mesh, nucleoplasm, and cytoplasm were quantified by RT-qPCR (left panel). The hygromycin sequence encoded in the constructs was used as an internal control for normalization. The nuclear RNA clearance index is defined as the ratio of Chr/RNP mesh to nucleoplasmic RNA levels (right panel).

To distinguish between altered RNA production and nuclear RNA clearance, we fractionated cells and quantified spike-in-normalized RNA in chromatin-associated (Chr/RNP mesh) and nucleoplasmic compartments, approximating chromatin-proximal and soluble nuclear RNA pools, respectively. Integrating these profiles with nascent transcription measurements identified three response classes (Fig. 3C). Clusters 2 and 3, comprising the majority of response genes, accumulated nucleoplasmic RNA despite reduced nascent transcription, whereas cluster 1 showed limited accumulation. Consistent with this pattern, CHX treatment reduced the RNA clearance index, defined as the ratio of chromatin-associated to nucleoplasmic RNA, across all clusters, with the strongest effects in clusters 2 and 3 (Fig. 3D). Genes in clusters 2 and 3 were generally longer, contained more exons, and exhibited higher baseline RNA clearance indices despite relatively lower steady-state RNA abundance (Fig. 3D i–iv). They also displayed more ill-spliced transcripts by long-read Nanopore sequencing (Fig. 3D v–vi), consistent with a greater reliance on RNA processing and surveillance.

We next directly assess RNA stability using EU pulse-chase experiments. CHX broadly stabilized transcripts across the transcriptome, including mRNAs, nuclear-enriched lincRNAs, intronic transcripts, PROMPTs, and enhancer RNAs (Fig. 3F and Supplementary Fig. 3A, B). Histone mRNAs were also stabilized, consistent with their translation-coupled decay pathway^27–30^ (Supplementary Fig. 3C, D); this specialized gene class is discussed separately below. To quantitatively assess RNA metabolism, we combined time-resolved EU-seq with kinetic modeling to estimate transcript-specific synthesis and decay rates^31,32^. Cells were treated with CHX and pulse-labeled with EU for 60, 40, 20, or 10 min, followed by cell-number-normalized EU-RNA-seq and nuclear RNA-seq (Fig. 3G). Because the shortest labeling window was 10 min, the magnitude of the observed changes is likely conservative, although relative comparisons between conditions remained robust. CHX treatment caused coordinated reductions in both RNA synthesis and decay, indicating broad stabilization of nuclear transcripts (Fig. 3G). Notably, synthesis and decay remained positively correlated (Supplementary Fig. 3E), suggesting that coupling between RNA production and turnover is largely preserved despite their global reduction. These results indicate that translational inhibition shifts RNA metabolism into a lower-flux state, characterized by reduced transcriptional output and impaired RNA clearance.

To determine whether RNA stability depends on translational competence at the transcript level, we generated β-globin reporters with disrupted translation initiation or coding potential (Supplementary Fig. 3F). Translation-deficient reporters exhibited reduced RNA abundance in both nuclear and cytoplasmic fractions, whereas chromatin-associated nascent RNA remained largely unchanged, consistent with increased nuclear RNA clearance (Fig. 3H). Thus, translational competence promotes RNA stability. While acute translational inhibition globally stabilizes many nuclear RNAs, productive translation can also stabilize individual transcripts, consistent with a functional link between translation and RNA surveillance.

### RNA surveillance factors redistribute upon translation arrest

To explore the mechanism linking impaired RNA turnover and productive elongation, we first assessed whether ribotoxic stress response (RSR) was engaged. RSR is rapidly activated via ZAKα-dependent p38 signaling in response to ribosome collision^33,34^. In wild-type ESCs, both CHX and ANS induced p38 phosphorylation within 15 min (Supplementary Fig. 4A, i). However, in p53-deficient ESCs, CHX failed to induce detectable p38 or EIF2α phosphorylation even after 60 min, whereas ANS robustly activated p38 signaling (Supplementary Fig. 4A, i,ii). Despite these differences, CHX triggered comparable transcriptional responses in p53-deficient ESCs (Fig. 1C), arguing that canonical p38/EIF2α signaling is unlikely to account for the early nuclear transcriptional response.

To investigate nuclear contributions independently of cytosolic stress pathways, we quantified nuclear and cytoplasmic protein abundance using compartment-resolved quantitative tandem-mass-tag (TMT) proteomics. Notably, the abundance of the Pol II complex remained largely unchanged in the nucleus (Supplementary Fig. 4B). In contrast, several short-lived pluripotency factors, including KLF2, KLF5, and NANOG, were rapidly depleted in the nucleus (Supplementary Fig. 4C), yet Pol II and TBP occupancy at the TSSs of their target genes remained largely intact (Supplementary Fig. 4D). Together with the stability of core transcription machinery and promoter occupancy, these findings suggest that early transcriptional defects are unlikely to arise from impaired transcription factor activity or defective promoter engagement.

Next, we asked whether translational arrest induces nucleocytoplasmic protein redistribution. Indeed, acute CHX treatment for 60 min induced widespread, albeit generally modest, nucleocytoplasmic redistribution (Fig. 4A and Supplementary Fig. 4E). Proteins shifting from the nucleus to the cytoplasm were strongly enriched for RNA catabolism, mRNA processing, snRNP assembly, and modestly enriched for Pol II elongation and translation pathways (Fig. 4B). In contrast, nuclear-accumulating proteins were associated with mRNA transport, Pol II initiation, and DNA replication (Fig. 4B). Notably, nearly half of cytoplasmic-accumulating proteins increased by >10% within 60 min and exhibited a corresponding decrease in nuclear abundance, consistent with reciprocal redistribution (194/452; Supplementary Fig. 4F). These patterns suggest that translational arrest preferentially redistributes RNA-processing and RNA catabolic machinery to the cytoplasm.

**Figure 4.**
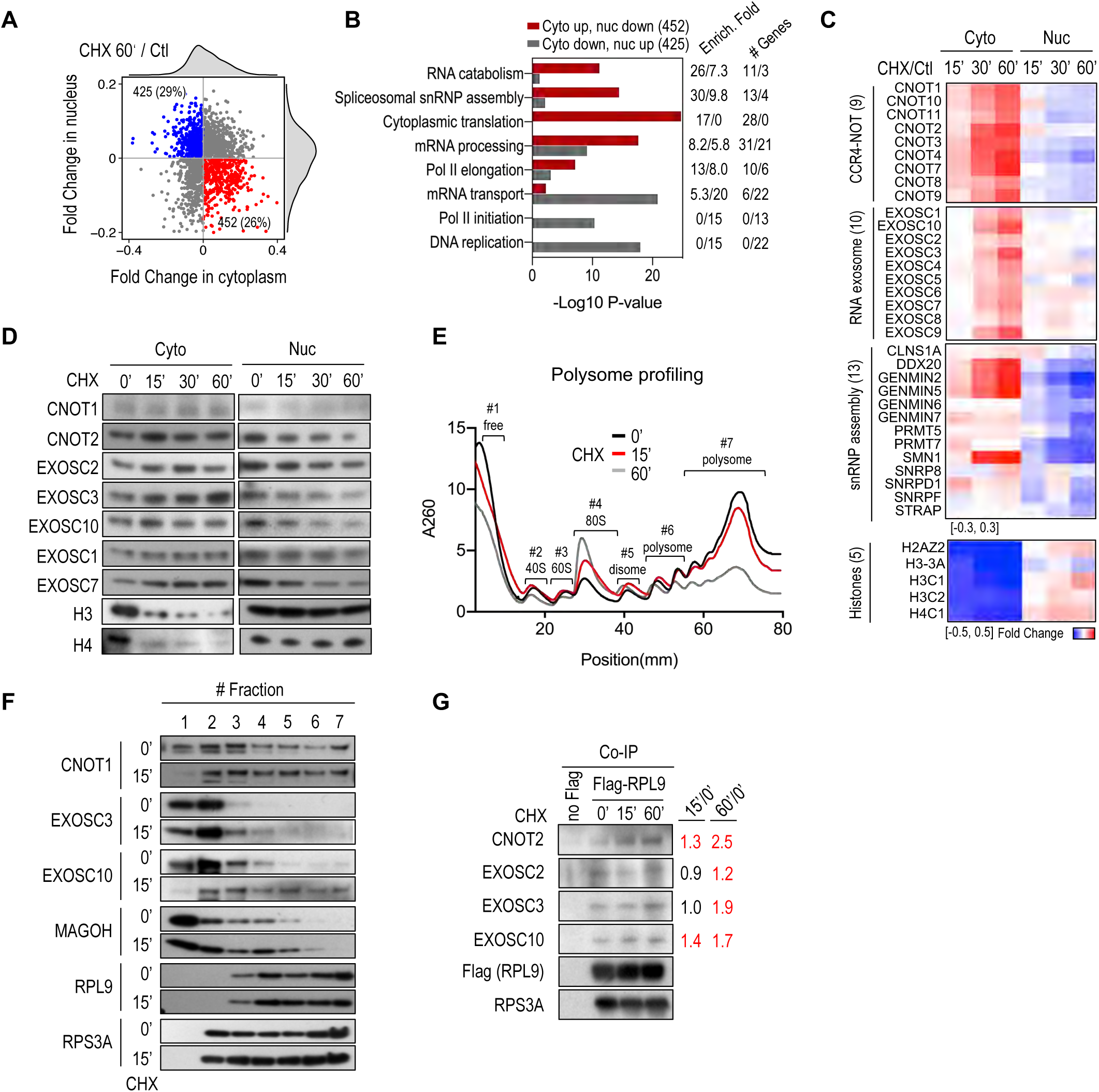
Redistribution of RNA surveillance factors upon translation arrest. (A) Dot plot showing nucleocytoplasmic protein redistribution following 60 min CHX treatment, based on quantitative mass spectrometry. Proteins with increased cytoplasmic and decreased nuclear abundance are shown in red (n = 452), whereas those with the opposite pattern are shown in blue (n = 425) (see also Supplementary Fig. 4E). (B) Gene Ontology (GO) enrichment analysis of proteins exhibiting reciprocal redistribution using the DAVID Functional Annotation Tool. Bars represent -log_10_ P values for enriched GO terms in proteins with increased cytoplasmic and decreased nuclear abundance (red), or decreased cytoplasmic and increased nuclear abundance (gray), following 60-min CHX treatment. (C) Heatmap showing fold changes in protein abundance relative to control, including components of the CCR4-NOT complex, RNA exosome, snRNP assembly factors, and histones. (D) Western blot validation of representative proteins exhibiting redistribution upon CHX treatment. Cytoplasmic (Cyto) fractions correspond to the soluble fraction obtained by 0.1% NP-40 extraction, whereas nuclear (Nuc) fractions represent the insoluble fraction after 0.2% NP-40 wash (see also Method). Quantitative band intensities are shown in Supplementary Fig. 4G. (E) Polysome profiling of control and CHX-treated cells. The x-axis indicates fraction position along the sucrose gradient, and the y-axis represents A260 absorbance. Position 0 corresponds to the top of the gradient. Seven fractions, including free, 40S, 60S, 80S, disome, light polysome, and heavy polysome, were defined based on the polysome profile. (F) Western blot analysis of representative proteins across sucrose gradient fractions corresponding to polysome profile in Fig. 4E. Quantitative band intensities are shown in Supplementary Fig. 4J. (G) Co-immunoprecipitation of RPL9 from cytoplasmic fractions in ESCs expressing endogenous Flag-tagged RPL9 (Flag-RPL9). Wild-type cells lacking the Flag tag were used as a negative control. Quantitative band intensities are shown on the right (see also Supplementary Fig. 4K).

Notably, components of RNA-surveillance machineries involved in RNA catabolism underwent subunit-wide shifts from the nucleus to the cytoplasm over 15–60 min of CHX treatment (Fig. 4C), including the CCR4–NOT complex^35^, a major deadenylation machinery, and the RNA exosome complex^36^, a 3’–5’ RNA exonuclease. Western blotting and immunofluorescence confirmed progressive cytoplasmic accumulation and corresponding nuclear depletion of representative subunits, including EXOSC1, EXOSC2, EXOSC3, EXOSC7, EXOSC10, CNOT1, CNOT2, as well as EJC components EIF4A3, and Y14 (Fig. 4D and Supplementary Fig. 4G-I). Notably, subunits involved in snRNP assembly also exhibited widespread redistribution, indicating a concomitant loss of nuclear RNA-processing capacity under translation inhibition (Fig. 4C). In parallel, cytoplasmic histone proteins were rapidly depleted within 15 min of CHX treatment (Fig. 4C, D), reflecting a high translation activity and rapid turnover of histone proteins in the cytoplasm.

Together, these observations suggest that ongoing translation is required to maintain the nuclear availability of RNA-surveillance and processing factors as well as the stability of highly labile protein pools. Upon translational arrest, disruption of this homeostatic balance may reduce nuclear RNA-surveillance and processing capacity, leading to impaired nuclear RNA metabolism and consequently reduced productive Pol II elongation.

### RNA surveillance complexes are retained on stalled ribosome-associated assemblies

To investigate how RNA surveillance factors become retained in the cytoplasm following translational arrest, we examined their association with ribosomes by polysome profiling. Under basal conditions, heavy polysomes predominated, consistent with active translation in ESCs (Fig. 4E). CHX treatment rapidly collapsed heavy polysomes and increased 80S monosomes, disomes, and higher-order ribosome complexes, consistent with elongation blockade and ribosome collision, beginning within 15 min and further by 60 min (Fig. 4E).

Despite the overall reduction in active polysomes, components of the CCR4-NOT complex and RNA exosome, including CNOT1, EXOSC3, and EXOSC10, were redistributed from free, ribosomal-subunit, and monosome fractions toward disome- and polysome-containing fractions following CHX treatment (Fig. 4F and Supplementary Fig. 4J). Consistently, free fractions were markedly depleted of CNOT1 and EXOSC10 and substantially reduced for EXOSC3, suggesting reduced recycling and increased retention on ribosome-associated complexes. A similar pattern was observed for the exon junction complex (EJC) component MAGOH, which accumulated in monosome-, disome-, and polysome-containing fractions following CHX treatment (Fig. 4F). Because EJC complexes are normally displaced during early rounds of translation^22^, their retention in ribosome-associated fractions suggests impaired translation-coupled mRNP remodeling and prolonged association of RNA regulatory factors with stalled ribosomes.

To directly test whether RNA-surveillance machinery physically associates with ribosomes under translational arrest, we performed co-immunoprecipitation (co-IP) using cells expressing endogenous Flag-tagged RPL9. Components of both the CCR4-NOT complex and RNA exosome, including CNOT2, EXOSC2, EXOSC3, and EXOSC10, specifically co-purified with ribosomes, and these interactions increased over 15–60 min of CHX treatment (Fig. 4G and Supplementary Fig. 4K). These results provide direct biochemical evidence that translational arrest promotes association of RNA surveillance machinery with ribosome-containing complexes. Together, these findings suggest that translational arrest not only redistributes RNA-surveillance factors to the cytoplasm but also triggers their retention on stalled ribosome-associated assemblies. Such retention would be expected to limit recycling of RNA-surveillance machinery and thereby reduce its nuclear availability.

### Loss of RNA surveillance phenocopies translation-dependent elongation defects

To test whether impaired RNA surveillance contributes functionally to transcriptional repression, we acutely depleted core components of two major RNA surveillance pathways: the RNA exosome subunits EXOSC2 and EXOSC3, and the EJC component Y14, using auxin-inducible degradation (AID) or dTAG systems (Fig. 5A). Cell-number-normalized EU-seq revealed rapid, genome-wide reductions in nascent transcription within 1-2 h of factor depletion (Fig. 5B-D and Supplementary Fig. 4L), indicating that RNA surveillance activity is required to sustain ongoing transcription.

**Figure 5.**
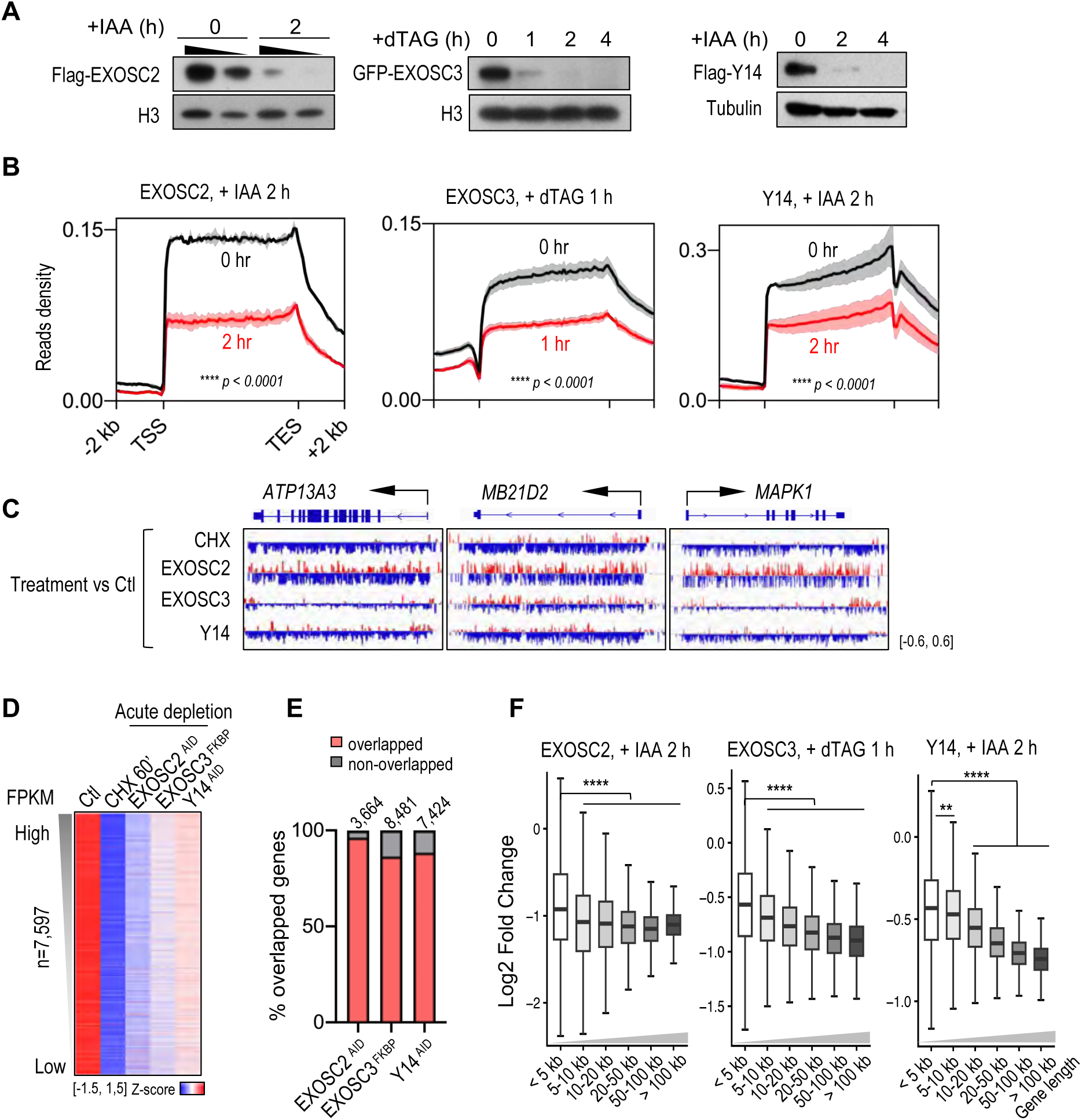
RNA surveillance loss phenocopies CHX effects on transcription. (A) Western blot validation of rapid degradation of EXOSC2, EXOSC3 and Y14. Anti-Flag antibodies were used for detection of EXOSC2 and Y14, and anti-GFP antibody was used for EXOSC3. IAA was used to induce degradation of AID-tagged EXOSC2 and Y14, whereas dTAG was used to degrade FKBP-tagged EXOSC3 in ESC. ESCs were treated with IAA (0.5 mM) or dTAG (1 *μ*M) for indicated times. (B) Metagene analysis of cell-number normalized EU-seq (10 min labeling) across 22,470 mouse protein-coding gene bodies with ±2 kb flanking regions. Only the sense strand is shown. Solid lines indicate mean of two biological replicates and shaded areas represent SEM. Statistical significance was assessed using a two-sided Kolmogorov-Smirnov test. Spike-in controls were used for normalization. (C) Representative IGV tracks showing fold changes in EU-seq signal at the *ATP13A3*, *MB21D2*, and *MAPK1* loci under CHX treatment or upon depletion of EXOSC2, EXOSC3, and Y14. Tracks represent merged biological replicates. (D) Heatmap showing cell-number normalized gene expression changes following 60 min CHX treatment or acute degradation of EXOSC2, EXOSC3, or Y14 relative to control cells within each batch. Genes with FPKM > 1 in control cells were included and ranked by expression level. Values are shown as Z-scores. (E) Overlap analysis of altered genes (p < 0.05) upon acute protein degradation and after 60 min CHX treatment. Total gene numbers per condition are indicated above bars. (F) Boxplots showing EU-seq signal changes upon acute protein degradation stratified by gene length. Genes with FPKM > 1 were grouped into six categories based on gene length (<5, 5-10, 10-20, 20-50, 50-100, and >100 kb). Gene numbers per group are listed for EXOSC2, EXOSC3, and Y14 depletion, respectively: 1033/1064/1772/2654/1388/1033; 1496/1186/1859/2703/1396/992; 959/1072/1713/2536/1317/957. Boxes indicate the 25^th^ -75^th^ percentiles with medians shown, and whiskers represent the 5^th^ -95^th^ percentiles. Statistical significance was assessed using a two-sided Wilcoxon rank-sum test. Statistical comparisons were made between each group and the first group (<5 kb).

Notably, more than 80% of genes significantly downregulated upon acute depletion of EXOSC2, EXOSC3, or Y14 overlapped with those repressed after 60 min of CHX treatment (Fig. 5E), revealing a strong transcriptome-level convergence between disruption of RNA surveillance and translational inhibition. Consistently, transcriptional repression induced by depletion of RNA surveillance factors was strongly gene-length dependent, with long genes showing preferential sensitivity (Fig. 5F), closely mirroring the response to translational inhibition (Fig. 1G and Supplementary Fig. 1G–H).

Although acute AID-mediated degradation produces near-complete loss of individual surveillance factors, translational inhibition causes partial redistribution of many RNA regulatory proteins. While any single redistribution event is modest, the collective effects of multiple surveillance and processing factors are likely substantial. The qualitative similarity between these perturbations suggests that disruption of RNA surveillance is sufficient to recapitulate a major feature of the transcriptional response to translational arrest. Together, these results establish that RNA surveillance and processing pathways are required to sustain productive transcriptional elongation, particularly across long genes. They also provide functional evidence that the elongation defects observed following translational inhibition arise, at least in part, from impaired nuclear RNA surveillance capacity.

### Histone genes escape translation-dependent elongation defects through initiation-driven transcription

The preceding results indicate that translational inhibition primarily compromises productive elongation, with long genes showing the greatest sensitivity. We therefore asked whether any transcriptional programs escape this elongation-dependent repression. Among the small subset of genes that increased transcription following translational inhibition, replication-dependent histone genes represented the most prominent exception. Histone genes are short, lack polyadenylation, and undergo specialized 3’-end processing, reducing their dependence on sustained Pol II processivity and canonical co-transcriptional RNA-processing pathways^37,38^. They were rapidly upregulated across multiple translation inhibitors (CHX, HAR, and ANS) in both wild-type and p53-deficient ESCs using 3- and 10-min EU labeling (Fig. 6A).

**Figure 6.**
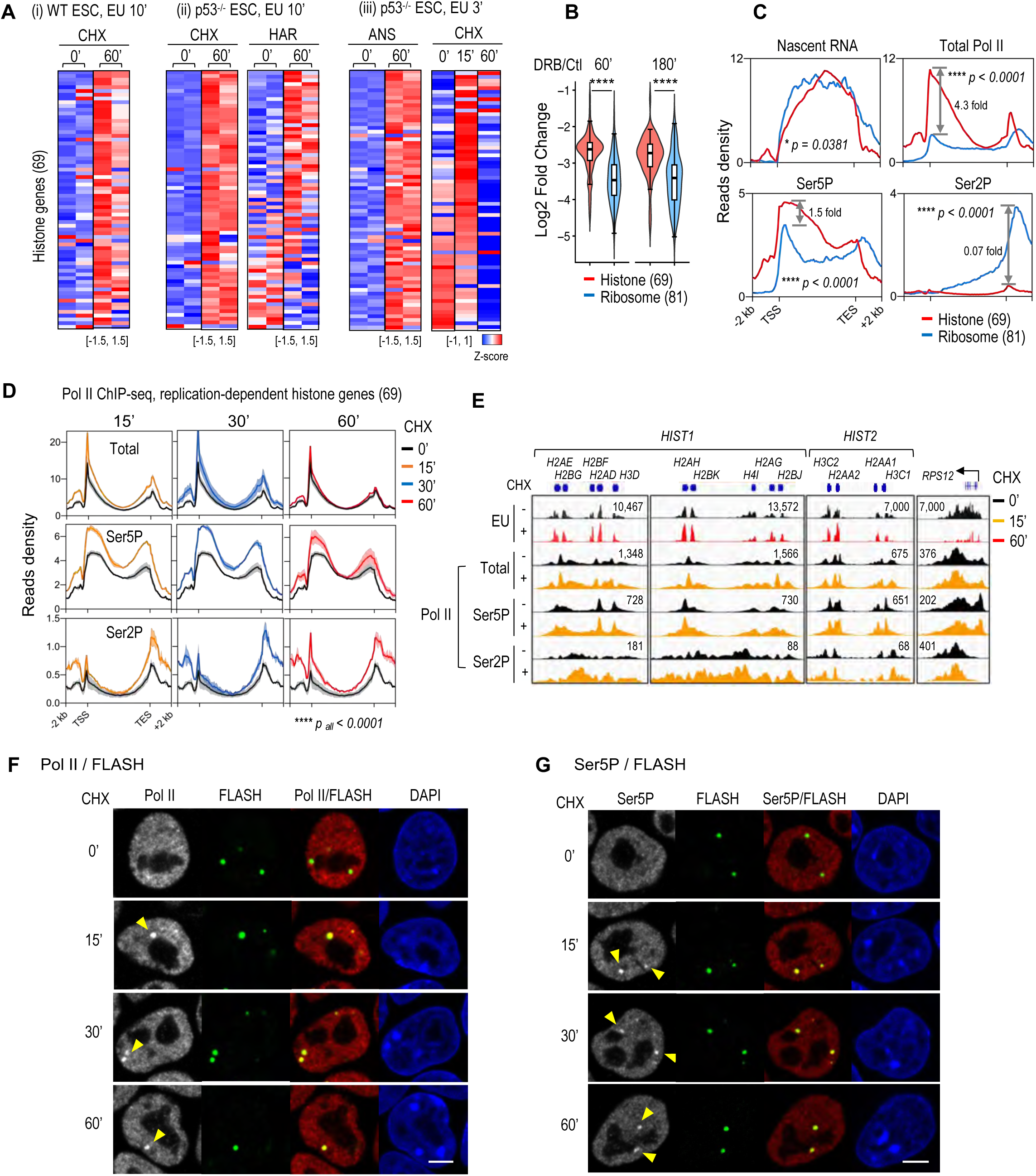
Transcriptional induction of replication-dependent histone genes upon translational inhibition. (A) Heatmap showing changes in EU-seq signal at replication-dependent histone genes following treatment with translational inhibitors in wild-type and p53^-/-^ ESCs at the indicated time points. Signals from two biological replicates are shown as Z-score-normalized values. (B) Violin plots showing changes in nascent transcription of histone genes and ribosomal protein genes following 60 and 180 min DRB treatment. Boxes indicate the 25^th^ -75^th^ percentiles with medians shown, and whiskers represent the 5^th^ -95^th^ percentiles. Outliers were omitted. Statistical significance was assessed using a two-sided Wilcoxon rank-sum test. (C) Metagene profiles of histone and ribosomal proteins genes. EU-seq (3 min) and Pol II occupancy (Total Pol II, Ser5P, and Ser2P) with ±2 kb flanking regions are shown. Fold differences between histone and ribosomal protein genes at indicated positions are shown in grey. Only the sense strand is shown. Statistical significance was assessed using a two-sided Kolmogorov-Smirnov test. Spike-in controls were used for normalization in EU-seq. (D) Metagene profiles of Total, Ser5P, and Ser2P Pol II ChIP-seq signals across 69 replication-dependent histone genes and ±2 kb flanking regions. Only the sense strand is shown. Solid lines indicate mean of two biological replicates and shaded areas represent SEM. Statistical significance was assessed using a two-sided Kolmogorov-Smirnov test. Signals were normalized by sequencing depth. (E) Representative IGV snapshots showing EU-seq (3 min) and Pol II ChIP-seq signals at histone clusters (*HIST1* and *HIST2*) and the *RPS12* locus. (F–G) Immunofluorescence showing CHX-induced redistribution of Pol II at histone loci and its spatial relationship with histone locus bodies marked by FLASH. Induced Pol II foci are indicated by yellow arrows. Representative cells are shown. Scale bars: 5 *μ*m.

To directly test their elongation dependence, we inhibited Pol II pause-release and productive elongation activity using DRB. While transcription of ribosomal protein genes was strongly suppressed, histone genes were less affected (Fig. 6B). Accordingly, compared with similarly highly expressed ribosomal protein genes, histone genes displayed markedly higher Total Pol II and Ser5P occupancy but much lower Ser2P occupancy (Fig. 6C), reinforcing their lower dependency of Pol II elongation.

Within 15 min of translational inhibition, Total Pol II and Ser5P chromatin occupancy increased further at histone genes, whereas Ser2P remained comparatively low although increased occupancy (Fig. 6D-E). Consistently, Total and Ser5P Pol II rapidly accumulated at histone locus bodies (HLBs), specialized nuclear compartments marked by FLASH^37–39^ (Fig. 6F, G and Supplementary Fig. 5A, B), whereas Ser2P Pol II showed limited enrichment (Supplementary Fig. 5C, D). These data suggest that histone gene activation is dominated by promoter-proximal Pol II recruitment and initiation-associated Ser5 phosphorylation rather than canonical Ser2P-associated elongation. In parallel, histone mRNAs accumulated across nuclear and cytoplasmic compartments, and EU pulse-chase experiments showed stabilization upon CHX treatment (Supplementary Fig. 3C-D).

Together, these observations indicate that histone transcription is predominantly initiation-driven, with CHX-induced activation reflecting enhanced promoter-proximal recruitment rather than Ser2P-dependent elongation. Histone gene output is further supported by translation-coupled RNA stabilization. Collectively, these features establish replication-dependent histone genes as a notable exception to the translation- and surveillance-dependent elongation defects affecting most protein-coding genes.

## Discussion

Gene expression is commonly viewed as a linear pathway from transcription to translation. Our findings reveal a reciprocal layer of regulation in which ongoing translation feeds back to sustain productive transcription elongation through maintenance of nuclear RNA-surveillance capacity (Fig. 7). Acute translational inhibition rapidly suppresses nascent transcription, impairs Pol II pause release and elongation, and reduces transcriptional output before substantial loss of chromatin-associated Pol II. At the same time, nuclear RNAs transiently accumulate owing to impaired RNA turnover. Together, these observations uncover a rapid cross-compartment feedback mechanism that links translational activity to transcriptional competence and nuclear RNA homeostasis.

**Figure 7.**
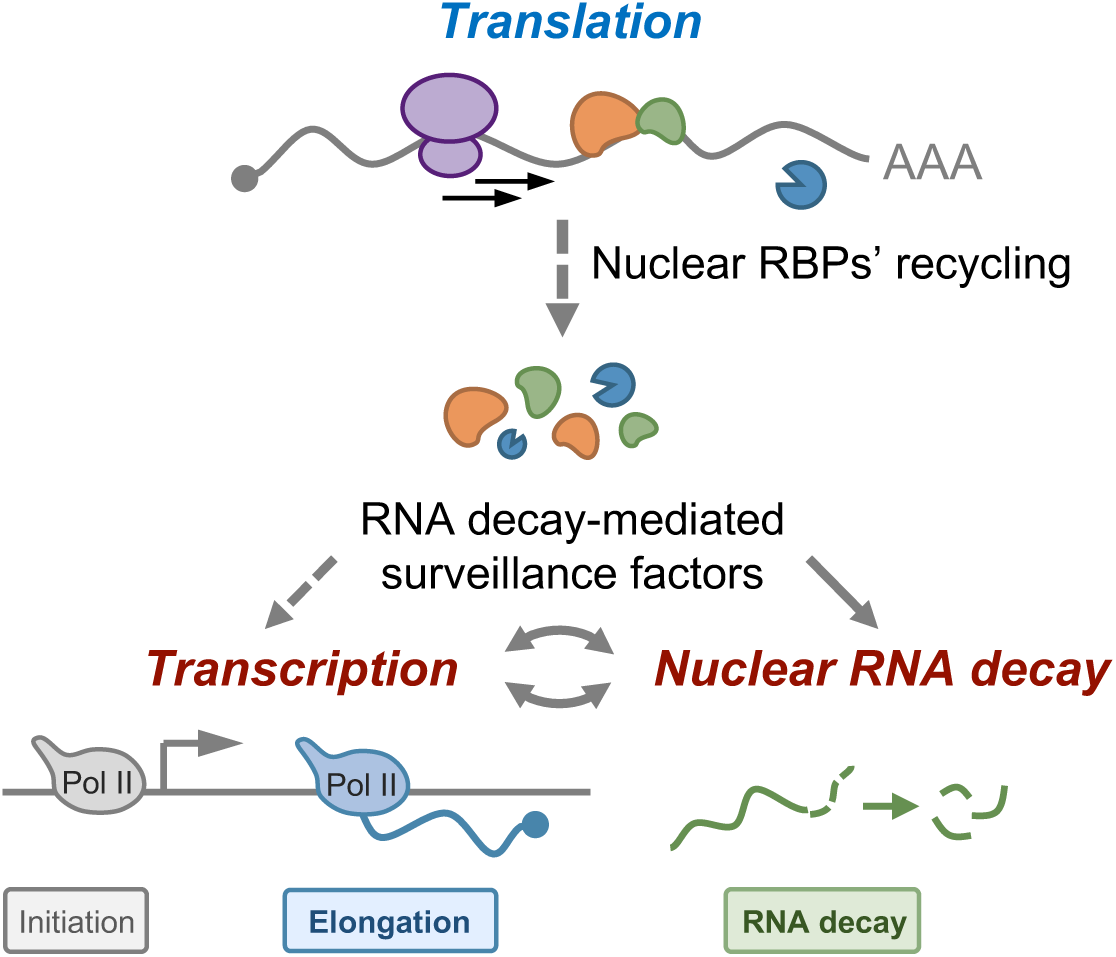
Model linking ongoing translation to nuclear RNA surveillance and transcriptional elongation. Ongoing translation sustains nuclear RNA-surveillance capacity by enabling continuous recycling of RNA-processing and RNA-surveillance factors between the cytoplasm and nucleus. This surveillance activity supports co-transcriptional RNA processing and productive RNA Pol II elongation, particularly at genes that require sustained processivity. Upon translation inhibition, RNA-surveillance factors become retained in stalled ribosome-associated cytoplasmic assemblies, reducing their nuclear availability. The resulting decline in nuclear RNA-surveillance capacity impairs RNA turnover and productive Pol II elongation, leading to reduced transcriptional output, particularly at long genes. Through this feedback mechanism, translational activity helps maintain transcriptional competence and coordinates RNA production with the cellular capacity for RNA processing and turnover.

A central conclusion of our study is that productive elongation is a dynamically maintained state rather than a passive consequence of Pol II chromatin occupancy. Following translational inhibition, engaged polymerases remain associated with chromatin while transcriptional output rapidly declines, demonstrating that Pol II occupancy and productive transcription can become uncoupled. The preferential sensitivity of long genes further supports this model. Long transcription units require sustained processivity, efficient pause release, co-transcriptional RNA processing, and continuous surveillance, making them particularly vulnerable to reductions in nuclear RNA-regulatory capacity. In contrast, transcription units that rely less heavily on sustained elongation are comparatively resistant. Consistent with this framework, replication-dependent histone genes represent a notable exception. Their transcription appears to rely predominantly on Pol II promoter-proximal recruitment and initiation-associated Ser5P rather than extensive Ser2P-dependent elongation. The relative resistance of histone genes to translation-induced elongation defects therefore reinforces the idea that gene-specific vulnerability is determined by dependence on productive elongation and co-transcriptional RNA processing.

Mechanistically, our data identify nuclear RNA-surveillance capacity as a critical link between translation and transcription. Multiple RNA-processing and surveillance machineries, including CCR4-NOT, the RNA exosome, and EJC components, undergo coordinated redistribution from the nucleus to the cytoplasm following translational arrest. These factors become increasingly associated with stalled ribosome-containing assemblies, suggesting that translational arrest limits their recycling and reduces their nuclear availability. Acute depletion of representative surveillance factors phenocopies the transcriptional defects induced by translation inhibition, particularly at long genes, providing functional support for a causal relationship between RNA-surveillance capacity and productive elongation. β-globin reporter experiments confirm that translation competency stabilizes individual coding transcripts when surveillance is intact, whereas global translation inhibition impairs surveillance capacity, transiently stabilizing otherwise unstable nuclear RNAs. Although translational inhibition likely affects multiple cellular processes simultaneously, our findings suggest that impaired nuclear RNA surveillance contributes substantially to the early elongation defects observed following translational arrest.

This mechanism is distinct from canonical ribotoxic stress signaling. Stalled ribosomes can activate ZAKα-dependent p38/JNK under some conditions^33,34^. However, the rapid transcriptional response described here occurs in the absence of detectable activation of the p38 or EIF2α pathways in p53-deficient ESCs and precedes major changes in promoter occupancy or core transcription machinery. These observations argue that the early transcriptional consequences of translational arrest primarily reflect alterations in RNA-regulatory capacity rather than secondary stress-response programs. More broadly, linking RNA-surveillance capacity to translation may provide a mechanism for coordinating nuclear RNA production with the cellular capacity to process, monitor, and utilize transcripts.

Our findings extend previous studies highlighting intimate connections among transcription, RNA processing, RNA decay, and translation. Chromatin-associated RNA-binding proteins such as WDR43^33^ and PSPC1^34^, RNA-surveillance complexes, and multifunctional regulators such as RPB4/7^40,41^, the CCR4-NOT complex^42–45^, and the RNA exosome^20^, have been implicated in coordinating gene expression across nuclear and cytoplasmic compartments. Likewise, accumulating evidence indicates that co-transcriptional RNA processing influences transcription elongation and determines the fate of nascent transcripts^15,46^. Moreover, co-transcriptional RNA modifications, including m^6^A^44,47–49^, pseudouridine^50^, and A-to-I editing^51–53^, are preferentially deposited on coding transcripts and influence downstream translational efficiency. Our study adds an upstream regulatory layer to this framework by showing that ongoing translation helps sustain the nuclear availability of RNA-surveillance machinery, thereby maintaining productive Pol II elongation. In this view, translation is not merely a downstream consumer of gene products but also an active participant in preserving transcriptional competence.

Together, our results support a model in which ongoing translation sustains productive Pol II elongation by maintaining nuclear RNA-surveillance capacity. Translational arrest promotes retention of RNA-regulatory factors on stalled ribosome-associated complexes, reducing their nuclear availability, slowing RNA turnover, and impairing productive elongation, particularly at genes with high processing demands. This feedback mechanism links translational capacity to transcriptional fidelity and RNA homeostasis, revealing an unexpected route through which translation influences nuclear gene expression. More broadly, disruption of this translation-RNA surveillance feedback may represent a previously unrecognized source of transcriptional stress in pathological states^20,36,54,55^.

### Limitations

Our study establishes a link between translation, nuclear RNA surveillance, and productive Pol II elongation, however, several aspects remain unresolved. Our conclusions are primarily based on pharmacological perturbation of translation; complementary genetic approaches would further strengthen the causal framework. In addition, β-globin reporters capture transcript-level effects but represent a limited subset of transgenes-derived transcripts, thereby leaving the general principles governing translation-dependent endogenous RNA regulation to be fully elucidated. Although our data support a model in which translational activity influences transcription, at least in part through redistribution of nuclear surveillance proteins and consequent changes in nuclear RNA processing and decay, the precise molecular pathways linking these processes remain to be defined, and additional regulatory pathways may also contribute to this translation-dependent control of transcription. Finally, whether this regulatory coupling extends across diverse cell types and physiological or stress conditions beyond pharmacologically induced acute translational inhibition remains an open question.

## AUTHOR CONTRIBUTIONS

X.S. conceived and supervised the study. M.H. performed most of the experiments and bioinformatic analyses. .H. generated the p53^-/-^, EXOSC3^FKBP^ and Y14^AID^ cell lines and performed Y14 EU-seq experiments. R.D. performed ChIP experiments for Ser5P Pol II and TBP. Y.L. constructed reporter plasmids and cell lines and performed RNA expression analyses. Y.Z. conducted computational analysis of RNA dynamics analysis. X.H. performed Nanopore sequencing data analysis. Z.Z. generated RPL9-FLAG-tagged cell lines. L.X. provided Nanopore RNA-seq data and performed EU-seq in EXOSC3^FKBP^ cells. X.S. and M.H. wrote the manuscript.

## ACKNOWLEDGMENTS

We thank members of the Shen laboratory for insightful discussions and suggestions. We acknowledge the Isotope Laboratory at the Center of Biomedical Analysis, Tsinghua University, for technical support. This work was supported by the Fundamental and Interdisciplinary Disciplines Breakthrough Plan of the Ministry of Education of China (JYB2025XDXM503), the NSFC Excellence Research Group Program (32588201), and the National Key R&D Program of China (2024YFA0916602), as well as the State Key Laboratory of Membrane Biology, the Beijing Advanced Innovation Center for Genomics, and the Center for Life Sciences.

## Declaration of generation AI and AI- assisted technologies in the writing process

During the preparation of this manuscript, the authors used ChatGPT (OpenAI) solely to assist with language editing. The authors reviewed, revised, and verified all AI-assisted text and take full responsibility for the content, accuracy, and integrity of the final manuscript.

## Methods

### Cell culture

Mouse embryonic stem cells (ESCs) were cultured on 0.1% gelatin-coated plates in high-glucose DMEM supplemented with 15% fetal bovine serum, GlutaMAX, penicillin-streptomycin, MEM non-essential amino acids, nucleoside mix, 0.1 mM 2-mercaptoethanol, and 1,000 U/mL recombinant leukemia inhibitory factor (LIF). Cells were maintained at 37°C in a humidified incubator with 5% CO₂. For acute translation inhibition, cells were treated with cycloheximide (CHX; Selleck, S7418), harringtonine (HAR; Selleck, S9063) and anisomycin (ANS; Selleck S7409) were added to the medium for indicated times. For transcription inhibition, cells were treated with DRB (100 *μ*M; Abcam, ab120939) or ActD (1 *μ*g/mL)for indicated times.

### Construction of endogenous tagged cell lines

Mouse embryonic stem cells (ESCs, 46C) were genetically engineered using CRISPR/Cas9-mediated genome editing to generate knockout and endogenous tagged cell lines.

For *Trp53* knockout (*Trp53*^-/-^) ESCs, two sgRNAs targeting coding regions of *Trp53* (sgRNA-1: CTGCTAGCTCAGAACAAATG; sgRNA-2: GTTACACTGAGAACCACTGT) were cloned into U6 promoter-driven sgRNA expression vectors. ESCs were co-transfected with both sgRNA plasmids and a Cas9 expression vector using Lipofectamine 3000 (Thermo Fisher Scientific), according to the manufacturer’s instructions. Transfected cells were selected with puromycin (1 *μ*g/mL) for 3-4 days, followed by isolation of single-cell-derived colonies. Genomic deletions at the *Trp53* locus were identified by PCR amplification across the targeted region and confirmed by Sanger sequencing. Loss of p53 protein expression was validated by immunoblotting.

For endogenous C-terminal knock-in or tagging, sgRNAs were designed to target genomic sequences adjacent to the stop codons of genes of interest. The sgRNA used for Y14 AID tagging (GTCGACAGAGGATTTATCAA) was employed to introduce an auxin-inducible degron (AID-3×Flag) cassette at the Y14 C-terminus. The sgRNA used for EXOSC3 tagging (GACTGGCAGAGAGTTGACAT) was used to insert an EGFP-FKBP fusion tag at the endogenous EXOSC3 locus. For RPL9, the sgRNA (AAGCAGTCAGCTGCAGACAT) was designed to target the C-terminal region, and an AID-Flag tag was inserted at the endogenous RPL9 stop codon. EXOSC2^AID^ cells were previously generated in our laboratory^20^.

Donor templates containing coding sequences flanked by homology arms were co-transfected with sgRNA and Cas9 expression plasmids into ESCs using Lipofectamine 3000 (Thermo Fisher Scientific). Antibiotic selection was applied where appropriate to enrich for edited populations, followed by plating at low density for clonal isolation. Individual colonies were expanded and screened by PCR across both 5’ and 3’ integration junctions to confirm site-specific genomic insertion. Correctly targeted clones were verified by Sanger sequencing of PCR amplicons spanning the edited loci. Protein expression and expected molecular weight shifts were validated by immunoblotting using tag-specific and/or gene-specific antibodies. All validated ESC lines were derived from independent clones and used for downstream analyses.

### Nascent peptide labeling and detection by Click chemistry

Global translation activity was assessed using O-propargyl-puromycin (OP-puro; Jena Bioscience, NU-931) or L-Homopropargyl Glycine (L-HPG; GLPBIO, GC14450), which incorporate into nascent polypeptide chains as a puromycin and methionine analogs, respectively.

For measuring translation output in intact cells, mouse embryonic stem cells (ESCs) were plated at a density of 1 × 10⁴ cells per well on 0.25% Matrigel-coated coverslips in 24-well plates at least 12 h prior to treatment. Cells were treated with IAA for the indicated duration, followed by incubation with 20 *μ*M OP-puro for 5 min at 37°C. For HPG labeling, medium was replaced by methionine (Met)-free medium, and cells were incubated for 30 minutes to reduce cellular Met pools. Then, 5 *μ*M L-HPG was added to Met-free medium and cells were labeled for the indicated times. After labeling, cells were washed with PBS and fixed with 4% paraformaldehyde for 15 min at room temperature. Cells were then permeabilized with 0.5% Triton X-100 for 10 min. Following a brief PBS rinse, cells were incubated with click reaction mixture in the dark at room temperature for 30 min. The click reaction mixture was prepared as follows: 50 mM HEPES pH 7.5, 5 mM Tris(3-hydroxypropyltriazolylmethyl)amine (THPTA; Sigma 762342), 2.5 mM CuSO₄, 0.1 mM Alexa Fluor 647 azide (Invitrogen, A10277), and 10 mM sodium ascorbate. Nuclei were stained with 1 *μ*g/mL Hoechst (diluted in PBS) for 10 min and cells were mounted with Fluoromount G for imaging.

For nuclear extraction experiments, HPG-labeled cells were incubated in hypotonic lysis buffer containing 0.1% NP-40 (20 mM HEPES pH 7.5, 10 mM KCl, 1.5 mM MgCl₂, 1 mM EDTA, 10% glycerol, 0.1 mM Na₃VO₄, 1 mM DTT, 1 mM PMSF, and 1× protease inhibitor cocktail). Cells were washed twice with the same buffer and then fixed with 4% paraformaldehyde for 15 min at room temperature. All subsequent steps, including permeabilization, click chemistry reaction, nuclear staining, and imaging, were performed as described above.

For OP-puro labeling in isolated nuclei, 5 × 10^6^ cells were washed twice with ice-cold PBS and resuspended in five volumes of hypotonic buffer A (20 mM HEPES pH 7.5, 10 mM KCl, 1.5 mM MgCl₂, 10% glycerol, 1 mM EDTA, 0.1 mM Na₃VO₄, 1 mM DTT, 1 mM PMSF, and protease inhibitors) supplemented with 0.2% NP-40. After incubation on ice for 5 min, nuclei were pelleted by centrifugation at 1,300 × g for 3 min at 4°C. The nuclear pellet was washed twice with freezing buffer (50 mM Tris-HCl pH 8.0, 5 mM MgCl₂, 0.1 mM EDTA, 40% glycerol) and resuspended in the same buffer. Nuclei were pre-treated with inhibitors as indicated for 20 min, followed by addition of an equal volume of 2× transcription buffer (final concentrations: 25 mM Tris-HCl pH 8.0, 150 mM KCl, 5 mM MgCl₂, 0.05 mM EDTA, 20% glycerol, 2 mM DTT, 1 mM each ATP, CTP, GTP, and UTP, and 20 *μ*M OP-puro). Reaction was carried out for 20 min at 30°C. Nuclei were then fixed with 4% paraformaldehyde and processed for click chemistry and imaging as described above.

### Immunofluorescence staining

Culture dishes were coated with 0.25% Matrigel and incubated at 37 °C for 2 h. Cells were seeded at 30-50% confluence and cultured overnight. Following experimental treatments, the culture medium was aspirated, and cells were rinsed once with PBS. Cells were fixed with 4% paraformaldehyde for 15 min and then permeabilized in PBS containing 3% BSA and 0.1% Triton X-100 for 30 min to 1 h at room temperature. After blocking, cells were incubated in primary antibody diluted in blocking solution or PBS for 1 h at room temperature or overnight at 4 °C. Following three washes with PBS, fluorophore-conjugated secondary antibody (diluted 1:1000 in PBS) was applied and incubated for 1 h at room temperature in the dark. Nuclei were stained with 1 *μ*g/mL Hoechst (diluted in PBS) for 10 min. After three final washes with PBS, excess liquid was removed, and coverslips were mounted with Fluoromount G (SouthernBiotech) and imaged on a Nikon A1R-HD multiphoton microscope. Imaging parameters were kept constant across conditions.

### Nascent RNA labeling by EU incorporation

Culture dish coating and cell seeding were performed as described for immunofluorescence staining. Prewarmed culture medium containing 2 mM 5-ethynyl uridine (EU) was added to cells to label nascent RNA for 10 min. Immediately after labeling, cells were fixed with 4% paraformaldehyde for 15 min at room temperature and permeabilized with PBS containing 0.1% Triton X-100 for 30 min. Following a brief PBS rinse, cells were incubated with click reaction mixture for 30 min in the dark at room temperature. The reaction mixture contained 50 mM HEPES pH 7.5, 5 mM THPTA, 2.5 mM CuSO₄, 0.1 mM Alexa Fluor 647 azide, and 10 mM sodium ascorbate. Nuclei were stained with 1 *μ*g/mL Hoechst for 10 min, and coverslips were mounted with Fluoromount-G for imaging.

### EU-seq

EU-seq was performed as previously described with modifications^20^. ESCs were metabolically labeled with 1 mM EU for 3- or 10-min. Cells were washed with PBS and were immediately lysed in 1 mL TRIzol. EU-labeled drosophila cells were added as spike-in controls prior to RNA extraction. Total RNA (10-20 *μ*g) was subjected to genomic DNA removal using RQ1 DNase I at 37 °C for 30 min, followed by RNA fragmentation at 94 °C for 5 min and purification by ethanol precipitation. For biotinylation of EU-labeled RNA, purified RNA was incubated in a 50 *μ*L reaction containing 50 mM HEPES pH 7.5, 2.5 mM THPTA, 2.5 mM CuSO₄, 4 mM Biotin-PEG3-azide (Aladdin, B122225), and 10 mM sodium ascorbate at room temperature for 1 h. The reaction was quenched with 450 *μ*L 5 mM EDTA, and RNA was purified via phenol-chloroform extraction followed by isopropanol precipitation and dissolved in 100 *μ*L nuclease-free water. Tandem purification of biotinylated EU-labeled RNA was carried out using Dynabeads C1 streptavidin magnetic beads. Beads were pre-washed twice with 1× B&W buffer (5 mM Tris-HCl pH 7.5, 0.5 mM EDTA, 1 M NaCl, 0.005% Tween-20), resuspended in 100 *μ*L 2× HSWB (1× HSWB: 10 mM Tris-HCl pH 7.5, 1 mM EDTA, 0.1 M NaCl, 0.01% Tween-20). Biotinylated RNA was denatured at 65°C for 10 min and incubated with blocked beads for 30 min at room temperature with rotation. Beads were washed twice with 10× HSWB at 50°C for 2 min, followed by two washes with nuclease-free water at 50°C with 1,200 rpm shaking. RNA was eluted twice in 50 *μ*L NLS buffer (50 mM Tris-HCl pH 8.1, 10 mM EDTA, 1% SDS) at 95°C for 5 min, and eluates were pooled. A second round of purification was performed using 10 *μ*L fresh C1 beads (pre-washed by NLS buffer) following the same protocol. Finally, beads with bound RNA were resuspended in 10 *μ*L nuclease-free water and subjected to cDNA synthesis and library construction with the Scale ssDNA-seq Lib Prep Kit (Abclonal, RK20222) or directly for strand-specific library preparation with the Fast RNA-seq Lib Prep Kit (Abclonal, RK20306).

For EU-seq analysis, adapter trimming and quality filtering were performed using Trim Galore with parameters -q 20, --phred33, --stringency 12, --length 20, -e 0.1, and --paired. Filtered paired-end reads were aligned to the mouse reference genome (mm10) using HISAT2 with default settings. Gene-level read counts were obtained using featureCounts by summarizing reads mapped to annotated exons, and expression levels were calculated as FPKM. Spike-in normalization was performed by mapping reads to the dm6 and the corresponding spike-in reference using Bowtie2. Absolute RNA abundance was defined as the ratio of mouse-mapped reads to spike-in reads and subsequently normalized to the mean of control samples within each fraction. Normalized reads were further partitioned into genic and intergenic fractions based on genomic annotation^15^. Genome-wide coverage tracks (bigWig) were generated using bamCoverage with spike-in normalization. Metagene profiles were generated using ngs.plot. Differential expression analysis was performed using DESeq2 based on spike-in-normalized counts. Statistical significance was assessed using adjusted P values (Benjamini-Hochberg correction).

Fragments per kilobase of exon model per million mapped reads (FPKM) was calculated using the following formula: FPKM = (read counts × 10⁹) / (gene length × total mm10-mapped reads), where read counts represent the number of reads mapped to annotated exonic regions. Reads per kilobase per million mapped reads (RPKM) was calculated using an analogous formula: RPKM = (read counts × 10⁹) / (gene length × total mapped reads) with the distinction that RPKM was computed using reads mapped across full gene bodies, whereas FPKM was restricted to exonic regions. Spike-in normalization was performed using dm6-derived spike-in reads. A normalization factor (F1) was defined as the ratio of spike-in-mapped reads to mm10-mapped reads, and subsequently scaled such that the corresponding ratio in control samples was set to 1. All FPKM values were corrected by multiplying by the F1 factor, yielding spike-in–normalized expression values.

### ChIP-seq

Chromatin immunoprecipitation followed by sequencing (ChIP-seq) was performed using an optimized protocol to profile genome-wide protein-DNA interactions. Cells were crosslinked with 1% formaldehyde for 10 min at room temperature, and quenched with 125 mM glycine for 5 min. Fixed cells were washed twice with ice-cold PBS, counted, and combined with 5% human 293T cells as a spike-in control.

For antibody pre-binding, antibodies against Pol II-NTD (CST 14958), Pol II Ser2P (CST 13499), Pol II Ser5P (Abcam ab5408), NELF-E (Proteintech 10705-1-AP), and CDK9 (CST 2316) were incubated with Protein G dynabeads (Thermo Fisher Scientific) in dilution buffer (16.7 mM Tris-HCl pH 8.1, 167 mM NaCl, 1.2 mM EDTA, 1.1% Triton X-100) for 2 h at 4 °C with rotation. For NELF-E and CDK9 immunoprecipitation, Protein G beads were pre-blocked with 2 mg/mL BSA in dilution buffer overnight at 4 °C with rotation prior to antibody coupling. Antibody-bound beads were then washed three times with dilution buffer at 4 °C. For each biological replicate, 3 × 10⁶ cells were lysed in 200 *μ*L nuclear lysis buffer (50 mM Tris-HCl pH 8.1, 10 mM EDTA, 1% SDS, 1 mM PMSF, 1 mM DTT, 1/100 protease inhibitor cocktail) and sonicated at 25% power for 1 min 15 s (10 s on, 20 s off). The lysate was centrifuged at 15,000 rpm for 10 min at 4 °C, and the supernatant was collected.

Immunoprecipitation was performed by mixing 36 *μ*L supernatant with 144 *μ*L dilution buffer with antibody-bound beads and incubating at 4 °C for 4 h to overnight with rotation. Beads were subsequently washed at 4 °C for 5-10 min each with the following buffers: low-salt IP wash buffer (20 mM Tris-HCl pH 8.1, 150 mM NaCl, 1 mM EDTA, 1% Triton X-100, 0.1% SDS, 0.1% sodium deoxycholate), high-salt IP wash buffer (20 mM Tris-HCl pH 8.1, 500 mM NaCl, 1 mM EDTA, 1% Triton X-100, 0.1% SDS, 0.1% sodium deoxycholate), LiCl IP wash buffer (10 mM Tris-HCl pH 8.1, 250 mM LiCl, 1 mM EDTA, 0.5% NP-40, 0.5% sodium deoxycholate), and TE buffer (10 mM Tris-HCl pH 8.1, 1 mM EDTA). After an additional wash with 10 mM Tris-HCl pH 7.5, the beads were resuspended in 10 *μ*L ultrapure water.

Bead-bound chromatin was subjected to tagmentation using the TruePrep DNA Library Prep Kit V2 (Vazyme) at 37 °C for 10 min. The reaction was quenched by addition of low-salt wash buffer, followed by sequential washes with TE buffer and 10 mM Tris-HCl pH 8.1. Chromatin was then resuspended in 10 *μ*L lysis buffer (20 mM Tris-HCl, pH 7.5, 0.05% NP-40, and 0.1 mg/mL Proteinase K) and incubated at 65 °C for 4 h to reverse crosslinks and digest proteins. Proteinase K was subsequently inactivated by heating at 85 °C for 15 min. Tagmented DNA was directly amplified by PCR using indexed primers from the TruePrep Index Kit V2 (Vazyme). Final libraries were purified using AMPure XP beads according to the manufacturer’s instructions.

For ChIP-seq data analysis, sequencing adapters and low-quality bases were removed using Trim Galore. Filtered reads were aligned to the mouse reference genome (mm10) using Bowtie2. PCR duplicates were removed with Picard, and only uniquely mapped reads were retained using SAMtools. Genome-wide coverage tracks (BigWig) were generated with bamCoverage. Metagene profiling and heatmap visualization were performed using ngs.plot. Peak calling was carried out using MACS with default parameters. Comparative heatmaps between control and treatment conditions were generated using deepTools (computeMatrix) based on bigWig files.

Public ChIP-seq datasets for KLF5 (GSE49848) and NANOG (GSE137193) were obtained from GEO datasets.

### Subcellular fractionation for RNA detection

Nuclear-cytoplasmic fractionation was performed based on the method described with modifications^9^. For each sample, 5 × 10⁶ cells at approximately 80% confluence were harvested by trypsinization, washed once with ice-cold PBS, and pelleted by centrifugation. All subsequent steps were carried out at 4 °C or on ice. The cell pellet was resuspended in 200 *μ*L lysis buffer (10 mM Tris-HCl pH 7.5, 150 mM NaCl, 0.05%–0.1% NP-40, 1 mM DTT, 1/100 protease inhibitor cocktail, 1 mM PMSF) and incubated on ice for 5 min. The lysate was then carefully layered onto 500 *μ*L of high-sucrose cushion buffer (10 mM Tris-HCl pH 7.5, 150 mM NaCl, 24% sucrose) and centrifuged at 12,000 rpm for 10 min at 4 °C. The upper 700 *μ*L was collected as the cytoplasmic fraction. The nuclear pellet was resuspended in 200 *μ*L glycerol buffer (20 mM Tris-HCl pH 8.0, 75 mM NaCl, 0.5 mM EDTA, 50% glycerol, 1 mM DTT), and 200 *μ*L high-salt nuclear lysis buffer (20 mM HEPES pH 7.5, 300 mM NaCl, 7.5 mM MgCl₂, 0.2 mM EDTA, 1% NP-40, 1-3 M urea, 1 mM DTT) was added. The mixture was vortexed vigorously until a stringy chromatin precipitate formed, then centrifuged at 15,000 rpm for 2 min at 4 °C. The supernatant was collected as the nucleoplasmic fraction, and the pellet was retained as the insoluble nuclear fraction containing chromatin and associated RNA networks.

One-tenth of the cytoplasmic fraction, one-fifth of the nucleoplasmic fraction, and the entire insoluble nuclear fraction were each added to 1 mL TRIzol reagent (Thermo Fisher Scientific). ERCC spike-in RNA (200–500 ng) was added to each fraction, and the samples were heated at 65 °C with shaking for 10 min to ensure complete dissolution. RNA was extracted using the TRIzol reagent according to standard protocols. rRNA was depleted using the rRNA depletion module (Abclonal, RK20348), and strand-specific RNA-seq libraries were prepared with the Fast RNA-seq Lib Prep Kit (Abclonal, RK20306) using random primers for first-strand synthesis.

RNA-seq data processing followed the same pipeline as EU-seq analysis, with modifications in spike-in normalization. ERCC spike-in RNAs were added to RNA samples obtained from subcellular fractionation and used for normalization of expression levels across fractions.

### RNA dynamic detection

To quantitatively assess RNA metabolic dynamics, we combined time-resolved EU labeling with kinetic modeling to infer RNA synthesis, processing, and decay rates. Cells were treated with CHX for 60 min, followed by EU pulse labeling for 10, 20, 40, or 60 min within the final 60-min treatment window. All samples were collected at the 60-min time point and processed for spike-in-normalized EU-RNA-seq and ribo-minus nuclear RNA-seq. EU-labeled nascent RNA and nuclear RNA were processed in parallel to provide dynamic and steady-state RNA measurements.

RNA kinetic parameters were inferred using a count-based INSPEcT workflow implemented in R^31^. The analysis used the INSPEcT, BiocParallel, and TxDb.Mmusculus. UCSC.mm10.knownGene packages, and exon and intron reads were quantified at the gene level based on the mm10 annotation. Samples were analyzed separately for the Ctl and CHX conditions. EU-labeled samples included four labeling time points (10, 20, 40, and 60 min), whereas nuclear RNA samples were used as the mature RNA reference. Raw sequencing reads were first aligned to the mm10 genome using HISAT2 to generate BAM files for expression quantification, while Bowtie2 alignments against ERCC or dm6 reference sequences were used to derive spike-in normalization factors. Gene-level quantification was then performed using quantifyExpressionsFromBAMs with the parameters by = “gene”, allowMultiOverlap = FALSE, strandSpecific = 2, isPairedEnd = TRUE, and DESeq2 = TRUE. EU samples were summarized using a replicate-aware time-point design, whereas total samples were treated as a single mature RNA group. Precomputed spike-in normalization factors were applied to each sample and averaged within the EU and total groups for each condition, and these group-level factors were then used for column-wise normalization of exon/intron expression and variance matrices. Common genes shared by the nascent and mature exon/intron expression and variance matrices were retained to construct the nascentExpressions and matureExpressions input objects. INSPEcT was run with tpts = c(1/6, 1/3, 2/3, 1), labeling_time = 1/6, preexisting = FALSE, and degDuringPulse = FALSE, followed by kinetic fitting using modelRates(seed = 1). Synthesis, processing, degradation, total RNA, pre-mRNA, and rate-associated P values were exported from the fitted model.

### EU pulse-chase assay

Mouse ESCs were seeded at 30–50% confluency and incubated overnight in culture medium supplemented with 100 *μ*M EU for metabolic labeling. After 16 h, the EU-containing medium was removed, and cells were washed twice with pre-warmed (37°C) PBS to eliminate residual EU. Cells were then cultured in fresh medium for 1 h to allow clearance of intracellular EU.

For the chase experiment, cells were subsequently switched to fresh medium containing either DMSO or CHX, and harvested at the indicated chase time points. Cells were lysed using TRIzol reagent for RNA extraction. EU-labeled RNA was then purified following a standard EU-seq protocol, and sequencing libraries were prepared for transcriptome analysis.

RNA-seq data processing followed the same pipeline as EU-seq analysis, with modifications in normalization. rRNA read counts were used for between-sample normalization of expression levels across conditions.

### Subcellular fractionation for protein detection

Cells (5 × 10⁶) were washed twice with ice-cold PBS and resuspended in five volumes of hypotonic buffer A (20 mM HEPES pH 7.5, 10 mM KCl, 1.5 mM MgCl₂, 10% glycerol, 1 mM EDTA, 0.1 mM Na₃VO₄, 1 mM DTT, 1 mM PMSF, and 1/100 protease inhibitor cocktail) supplemented with 0.1–0.2% NP-40. After incubation on ice for 5 min, cells were centrifuged at 1,300 × g for 3 min at 4 °C. The supernatant was collected as the cytoplasmic fraction, and the pellet was retained as the nuclear fraction. Nuclear pellets were washed twice with hypotonic buffer A.

For TMT-based protein quantification, cells were lysed in hypotonic buffer A containing 0.2% NP-40. Nuclear pellets were resuspended in 8 M urea in PBS and sonicated on ice (9 cycles of 3 s on / 5 s off at 20% power). After centrifugation at 15,000 rpm for 10 min at 4 °C, the supernatant was collected as the nuclear protein extract. Both cytoplasmic and nuclear protein extracts were precipitated with 5 volumes of ice-cold acetone at −80 °C overnight. Precipitated proteins were pelleted by centrifugation at 15,000 rpm for 10 min at 4 °C and washed twice with ice-cold acetone. Protein pellets were air-dried briefly and resuspended in 8 M urea in PBS. Protein concentrations from each fraction were determined, and equal amounts were labeled using the TMTsixplex kit (Thermo Fisher Scientific). Labeled samples were pooled in equal amounts and analyzed by mass spectrometry for multiplexed quantitative proteomics.

For western blot analysis, cells were lysed in hypotonic buffer A containing 0.1% NP-40. Following centrifugation, the supernatant was collected as the cytoplasmic fraction. Nuclear pellets were washed once with hypotonic buffer A containing 0.2% NP-40. Both nuclear and cytoplasmic fractions were denatured in 1× SDS loading buffer (50 mM Tris-HCl pH 6.8, 2% SDS, 10% glycerol, 0.1% bromophenol blue, and 1% β-mercaptoethanol) at 95 °C for 5 min.

### Polysome Profiling

Polysome profiling was performed using sucrose density gradient centrifugation. A 10× sucrose gradient buffer (200 mM Tris-HCl pH 7.5, 1 M KCl, 50 mM MgCl₂) was prepared and used to make 10% (w/v) and 50% (w/v) sucrose working solutions, and CHX and DTT were added to the sucrose solutions to final concentrations of 100 *μ*g/mL and 1 mM, respectively. Cell lysis buffer consisted of 1× sucrose gradient buffer containing 0.5% Triton X-100, supplemented immediately before use with 1 mM DTT, 1× protease inhibitor cocktail, 100 *μ*g/mL CHX, and 0.1 mM PMSF. Cells were harvested by trypsinization in the presence of 100 *μ*g/mL CHX, washed once with PBS containing CHX, and resuspended in 450 *μ*L lysis buffer. Lysis was carried out for 5 min at 4 °C with rotation, and the lysate was centrifuged at 400g for 5 min at 4 °C to collect the supernatant. A 10%–50% sucrose gradient was prepared by first adding 6 mL of 10% sucrose solution to a centrifuge tube, followed by slowly underlaying approximately 5 mL of 50% sucrose solution using a syringe. The gradient was generated using a gradient station with the program “long, 10%–50%, S1/1,1:50/80.0,0/21”. For ultracentrifugation, 400 *μ*L of clarified lysate was carefully layered on top of the gradient. Sample pairs were balanced to within 0.01 g using lysis buffer and loaded into the SW41-Ti rotor in positions 1-4, 2-5, and 3-6. Centrifugation was performed at 41,000 rpm for 2 h at 4 °C. After centrifugation, the gradient fractionation system was flushed five times with nuclease-treated water. The system was configured with the SW41-Ti rotor program, “Suc 10%–50%” gradient settings, a collection volume of 0.5 mL per fraction, and A₂₆₀ monitoring. After installing the collection tubes, the baseline was zeroed three times (fluctuation < 0.004), and fractionation was initiated. All steps were performed on ice to maintain ribosome integrity.

### Co-immunoprecipitation (Co-IP) of RPL9

Cells (5 × 10⁷ per condition) were harvested by trypsinization and resuspended in 500 *μ*L lysis buffer (20 mM Tris-HCl pH 7.5, 100 mM KCl, 5 mM MgCl₂, 0.5% Triton X-100, 1 mM PMSF, 1 mM DTT, 0.1 mM Na₃VO₄, 1× protease inhibitor cocktail). After rotation at 4 °C for 5 min, the supernatant were collected by centrifugation at 400 g for 3 min at 4 °C, supplemented with KCl to a final concentration of 200 mM, and clarified by centrifugation at 16,000 g for 5 min at 4 °C. The supernatant was collected as the cytoplasmic lysates. For immunoprecipitation, anti-Flag M2 Dynabeads (100 *μ*L per sample) were washed twice with lysis-200 buffer (20 mM Tris-HCl pH 7.5, 200 mM KCl, 5 mM MgCl₂, 0.5% Triton X-100) without protease/phosphatase inhibitors. The cytoplasmic lysate was incubated with the washed beads at 4 °C for 2–4 h. Beads were then washed three times with 500 *μ*L lysis-200 buffer supplemented with 1× protease/phosphatase inhibitor cocktail and RNase inhibitor at 4 °C. Finally, beads were resuspended in 1× SDS loading buffer and proteins were denatured at 95 °C for 5 min.

### Nuclear run-on transcription assay

Cells were seeded one day prior to the experiment at 70-80% confluency unless otherwise indicated. For translation inhibition experiments, cells were pretreated with CHX for 15-60 min prior to harvest. At the end of treatment, cells were harvested and permeabilized in ice-cold permeabilization buffer (10 mM Tris-HCl pH 7.4, 300 mM sucrose, 10 mM KCl, 5 mM MgCl₂, 1 mM EGTA, 0.05% Tween-20, 0.1% NP-40, 0.5 mM DTT, protease inhibitors, and 4 U/mL RNase inhibitor) for 5 min on ice. Nuclei were collected by centrifugation at 1,000 × g for 5 min at 4°C, washed twice, and resuspended in nuclear storage buffer (10 mM Tris-HCl pH 8.0, 25% glycerol, 5 mM MgCl₂, 0.1 mM EDTA, 5 mM DTT, and RNase inhibitor) at a density of 1 × 10⁷ nuclei per 100 *μ*L. For nuclear run-on transcription assays, equal volumes of nuclei suspension and pre-warmed 2× run-on buffer (300 mM KCl, 1% Sarkosyl, 5 mM MgCl₂, 1 mM DTT, 250 *μ*M ATP/GTP/CTP, 0.5 *μ*L α-³²P-UTP and RNase inhibitor) were mixed and incubated at 37°C for 3 min to allow elongation of nascent transcripts. Reactions were terminated by addition of 5 volumes of TRIzol LS (Thermo Fisher, 10296028CN), followed by RNA extraction according to the manufacturer’s instructions. RNA products were analyzed by 2% agarose gel electrophoresis and visualized by autoradiography.

### Design and construction of *β*-globin reporter and cell lines

The β-globin construct and its corresponding mutant sequence were generously provided by the Professor Hong Cheng as previously described^56^. Briefly, the sequences were amplified by PCR from the original plasmids and subcloned into a PiggyBac (PB) transposon-based vector backbone containing a hygromycin resistance cassette and a PB transposase recognition element, generating PB-β-globin reporter constructs (PB-bG and PB-bG-mut). For stable integration, mouse embryonic stem cells (ESCs) were co-transfected with the PB reporter plasmids and a PiggyBac transposase expression vector using Lipofectamine 3000 (Thermo Fisher Scientific), according to the manufacturer’s instructions. At 24-48 h post-transfection, cells were selected with hygromycin to establish stable cell lines. Stable ESC clones were expanded and subsequently used for downstream subcellular fractionation experiments as described in the corresponding sections.

## References

1. Bulut-Karslioglu, A., Macrae, T.A., Oses-Prieto, J.A., Covarrubias, S., Percharde, M., Ku, G., Diaz, A., McManus, M.T., Burlingame, A.L., and Ramalho-Santos, M. (2018). The transcriptionally permissive chromatin state of embryonic stem cells Is acutely tuned to translational output. Cell Stem Cell 22, 369–383 e368. 10.1016/j.stem.2018.02.004.

2. Rosa-Mercado, N.A., Buskirk, A.R., and Green, R. (2024). Translation elongation inhibitors stabilize select short-lived transcripts. Rna 30, 1572–1585. 10.1261/rna.080138.124.

3. Sun, M., Schwalb, B., Pirkl, N., Maier, K.C., Schenk, A., Failmezger, H., Tresch, A., and Cramer, P. (2013). Global analysis of eukaryotic mRNA degradation reveals Xrn1-dependent buffering of transcript levels. Mol Cell 52, 52–62. 10.1016/j.molcel.2013.09.010.

4. Cheng, Z., and Brar, G.A. (2019). Global translation inhibition yields condition-dependent de-repression of ribosome biogenesis mRNAs. Nucleic Acids Res 47, 5061–5073. 10.1093/nar/gkz231.

5. Schwanhausser, B., Busse, D., Li, N., Dittmar, G., Schuchhardt, J., Wolf, J., Chen, W., and Selbach, M. (2011). Global quantification of mammalian gene expression control. Nature 473, 337–342. 10.1038/nature10098.

6. Cramer, P. (2019). Organization and regulation of gene transcription. Nature 573, 45–54. 10.1038/s41586-019-1517-4.

7. Bentley, D.L. (2014). Coupling mRNA processing with transcription in time and space. Nat Rev Genet 15, 163–175. 10.1038/nrg3662.

8. Hsin, J.P., and Manley, J.L. (2012). The RNA polymerase II CTD coordinates transcription and RNA processing. Genes Dev 26, 2119–2137. 10.1101/gad.200303.112.

9. Yin, Y., Lu, J.Y., Zhang, X., Shao, W., Xu, Y., Li, P., Hong, Y., Cui, L., Shan, G., Tian, B., et al. (2020). U1 snRNP regulates chromatin retention of noncoding RNAs. Nature 580, 147–150. 10.1038/s41586-020-2105-3.

10. Leader, Y., Lev Maor, G., Sorek, M., Shayevitch, R., Hussein, M., Hameiri, O., Tammer, L., Zonszain, J., Keydar, I., Hollander, D., et al. (2021). The upstream 5’ splice site remains associated to the transcription machinery during intron synthesis. Nat Commun 12, 4545. 10.1038/s41467-021-24774-6.

11. Berg, M.G., Singh, L.N., Younis, I., Liu, Q., Pinto, A.M., Kaida, D., Zhang, Z., Cho, S., Sherrill-Mix, S., Wan, L., and Dreyfuss, G. (2012). U1 snRNP determines mRNA length and regulates isoform expression. Cell 150, 53–64. 10.1016/j.cell.2012.05.029.

12. Shao, W., Bi, X., Pan, Y., Gao, B., Wu, J., Yin, Y., Liu, Z., Peng, M., Zhang, W., Jiang, X., et al. (2022). Phase separation of RNA-binding protein promotes polymerase binding and transcription. Nat Chem Biol 18, 70–80. 10.1038/s41589-021-00904-5.

13. Skalska, L., Beltran-Nebot, M., Ule, J., and Jenner, R.G. (2017). Regulatory feedback from nascent RNA to chromatin and transcription. Nat Rev Mol Cell Biol 18, 331–337. 10.1038/nrm.2017.12.

14. Herzel, L., Ottoz, D.S.M., Alpert, T., and Neugebauer, K.M. (2017). Splicing and transcription touch base: co-transcriptional spliceosome assembly and function. Nat Rev Mol Cell Biol 18, 637–650. 10.1038/nrm.2017.63.

15. Xu, J., Li, X., Hao, X., Hu, X., Ma, S., Hong, Y., Zhang, J., Yan, D., Deng, H., Na, J., et al. (2024). Co-transcriptional RNA processing boosts zygotic gene activation. bioRxiv, 2024.2009.2014.613088. 10.1101/2024.09.14.613088.

16. Mimoso, C.A., and Adelman, K. (2023). U1 snRNP increases RNA Pol II elongation rate to enable synthesis of long genes. Mol Cell 83, 1264–1279 e1210. 10.1016/j.molcel.2023.03.002.

17. Caizzi, L., Monteiro-Martins, S., Schwalb, B., Lysakovskaia, K., Schmitzova, J., Sawicka, A., Chen, Y., Lidschreiber, M., and Cramer, P. (2021). Efficient RNA polymerase II pause release requires U2 snRNP function. Mol Cell 81, 1920–1934 e1929. 10.1016/j.molcel.2021.02.016.

18. Boddu, P.C., Gupta, A.K., Roy, R., De La Pena Avalos, B., Olazabal-Herrero, A., Neuenkirchen, N., Zimmer, J.T., Chandhok, N.S., King, D., Nannya, Y., et al. (2024). Transcription elongation defects link oncogenic SF3B1 mutations to targetable alterations in chromatin landscape. Mol Cell 84, 1475–1495 e1418. 10.1016/j.molcel.2024.02.032.

19. Bi, X., Xu, Y., Li, T., Li, X., Li, W., Shao, W., Wang, K., Zhan, G., Wu, Z., Liu, W., et al. (2019). RNA targets ribogenesis factor WDR43 to chromatin for transcription and pluripotency control. Mol Cell 75, 102–116.e109. 10.1016/j.molcel.2019.05.007.

20. Han, X., Xing, L., Hong, Y., Zhang, X., Hao, B., Lu, J.Y., Huang, M., Wang, Z., Ma, S., Zhan, G., et al. (2024). Nuclear RNA homeostasis promotes systems-level coordination of cell fate and senescence. Cell Stem Cell 31, 694–716.e611. 10.1016/j.stem.2024.03.015.

21. Ling, Y.H., Liang, C., Wang, S., and Wu, C. (2026). Live-cell single-molecule dynamics of eukaryotic RNA polymerase machineries. Science 391, eads0960. 10.1126/science.ads0960.

22. Boehm, V., and Gehring, N.H. (2016). Exon Junction Complexes: Supervising the Gene Expression Assembly Line. Trends Genet 32, 724–735. 10.1016/j.tig.2016.09.003.

23. Harper, N.W., Birdsall, G.A., Honeywell, M.E., Ward, K.M., Pai, A.A., and Lee, M.J. (2025). RNA Pol II inhibition activates cell death independently from the loss of transcription. Cell. 10.1016/j.cell.2025.07.034.

24. Kim, H.J., Kim, N.C., Wang, Y.D., Scarborough, E.A., Moore, J., Diaz, Z., MacLea, K.S., Freibaum, B., Li, S., Molliex, A., et al. (2013). Mutations in prion-like domains in hnRNPA2B1 and hnRNPA1 cause multisystem proteinopathy and ALS. Nature 495, 467–473. 10.1038/nature11922.

25. Habelhah, H., Shah, K., Huang, L., Ostareck-Lederer, A., Burlingame, A.L., Shokat, K.M., Hentze, M.W., and Ronai, Z. (2001). ERK phosphorylation drives cytoplasmic accumulation of hnRNP-K and inhibition of mRNA translation. Nat Cell Biol 3, 325–330. 10.1038/35060131.

26. Yang, Q., Guo, H., Li, H., Li, Z., Ni, F., Wen, Z., Liu, K., Kong, H., and Wei, W. (2025). The CXCL8/MAPK/hnRNP-K axis enables susceptibility to infection by EV-D68, rhinovirus, and influenza virus in vitro. Nat Commun 16, 1715. 10.1038/s41467-025-57094-0.

27. Stimac, E., Groppi, V.E., Jr., and Coffino, P. (1984). Inhibition of protein synthesis stabilizes histone mRNA. Mol Cell Biol 4, 2082–2090. 10.1128/mcb.4.10.2082-2090.1984.

28. Peltz, S.W., and Ross, J. (1987). Autogenous regulation of histone mRNA decay by histone proteins in a cell-free system. Mol Cell Biol 7, 4345–4356. 10.1128/mcb.7.12.4345-4356.1987.

29. McLaren, R.S., and Ross, J. (1993). Individual purified core and linker histones induce histone H4 mRNA destabilization in vitro. J Biol Chem 268, 14637–14644.

30. Kaygun, H., and Marzluff, W.F. (2005). Translation termination is involved in histone mRNA degradation when DNA replication is inhibited. Mol Cell Biol 25, 6879–6888. 10.1128/mcb.25.16.6879-6888.2005.

31. de Pretis, S., Kress, T., Morelli, M.J., Melloni, G.E., Riva, L., Amati, B., and Pelizzola, M. (2015). INSPEcT: a computational tool to infer mRNA synthesis, processing and degradation dynamics from RNA- and 4sU-seq time course experiments. Bioinformatics 31, 2829–2835. 10.1093/bioinformatics/btv288.

32. Mukherjee, N., Calviello, L., Hirsekorn, A., de Pretis, S., Pelizzola, M., and Ohler, U. (2017). Integrative classification of human coding and noncoding genes through RNA metabolism profiles. Nat Struct Mol Biol 24, 86–96. 10.1038/nsmb.3325.

33. Vind, A.C., Zhong, F.L., and Bekker-Jensen, S. (2025). Death by ribosome. Trends Cell Biol 35, 615–626. 10.1016/j.tcb.2024.10.013.

34. Vind, A.C., Genzor, A.V., and Bekker-Jensen, S. (2020). Ribosomal stress-surveillance: three pathways is a magic number. Nucleic Acids Res 48, 10648–10661. 10.1093/nar/gkaa757.

35. Collart, M.A. (2016). The Ccr4-Not complex is a key regulator of eukaryotic gene expression. Wiley Interdiscip Rev RNA 7, 438–454. 10.1002/wrna.1332.

36. Schmid, M., and Jensen, T.H. (2018). Controlling nuclear RNA levels. Nat Rev Genet 19, 518–529. 10.1038/s41576-018-0013-2.

37. Marzluff, W.F., Wagner, E.J., and Duronio, R.J. (2008). Metabolism and regulation of canonical histone mRNAs: life without a poly(A) tail. Nat Rev Genet 9, 843–854. 10.1038/nrg2438.

38. Marzluff, W.F., and Koreski, K.P. (2017). Birth and Death of Histone mRNAs. Trends Genet 33, 745–759. 10.1016/j.tig.2017.07.014.

39. Armstrong, C., Passanisi, V.J., Ashraf, H.M., and Spencer, S.L. (2023). Cyclin E/CDK2 and feedback from soluble histone protein regulate the S phase burst of histone biosynthesis. Cell Rep 42, 112768. 10.1016/j.celrep.2023.112768.

40. Goler-Baron, V., Selitrennik, M., Barkai, O., Haimovich, G., Lotan, R., and Choder, M. (2008). Transcription in the nucleus and mRNA decay in the cytoplasm are coupled processes. Genes Dev 22, 2022–2027. 10.1101/gad.473608.

41. Harel-Sharvit, L., Eldad, N., Haimovich, G., Barkai, O., Duek, L., and Choder, M. (2010). RNA polymerase II subunits link transcription and mRNA decay to translation. Cell 143, 552–563. 10.1016/j.cell.2010.10.033.

42. Ke, S., Pandya-Jones, A., Saito, Y., Fak, J.J., Vågbø, C.B., Geula, S., Hanna, J.H., Black, D.L., Darnell, J.E., Jr., and Darnell, R.B. (2017). m(6)A mRNA modifications are deposited in nascent pre-mRNA and are not required for splicing but do specify cytoplasmic turnover. Genes Dev 31, 990–1006. 10.1101/gad.301036.117.

43. Louloupi, A., Ntini, E., Conrad, T., and Ørom, U.A.V. (2018). Transient N-6-Methyladenosine Transcriptome Sequencing Reveals a Regulatory Role of m6A in Splicing Efficiency. Cell Rep 23, 3429–3437. 10.1016/j.celrep.2018.05.077.

44. Slobodin, B., Han, R., Calderone, V., Vrielink, J., Loayza-Puch, F., Elkon, R., and Agami, R. (2017). Transcription impacts the efficiency of mRNA translation via co-transcriptional N6-adenosine methylation. Cell 169, 326–337.e312. 10.1016/j.cell.2017.03.031.

45. Slobodin, B., Bahat, A., Sehrawat, U., Becker-Herman, S., Zuckerman, B., Weiss, A.N., Han, R., Elkon, R., Agami, R., Ulitsky, I., et al. (2020). Transcription dynamics regulate poly(A) tails and expression of the RNA degradation machinery to balance mRNA levels. Mol Cell 78, 434–444.e435. 10.1016/j.molcel.2020.03.022.

46. Ma, S., Hong, Y., Chen, J., Xu, J., and Shen, X. (2025). Single-cell nascent transcription reveals sparse genome usage and plasticity. Cell 188, 6873–6891.e6823. 10.1016/j.cell.2025.09.003.

47. Xiang, Y., Laurent, B., Hsu, C.H., Nachtergaele, S., Lu, Z., Sheng, W., Xu, C., Chen, H., Ouyang, J., Wang, S., et al. (2017). RNA m(6)A methylation regulates the ultraviolet-induced DNA damage response. Nature 543, 573–576. 10.1038/nature21671.

48. Meyer, K.D. (2019). DART-seq: an antibody-free method for global m(6)A detection. Nat Methods 16, 1275–1280. 10.1038/s41592-019-0570-0.

49. Powley, I.R., Kondrashov, A., Young, L.A., Dobbyn, H.C., Hill, K., Cannell, I.G., Stoneley, M., Kong, Y.W., Cotes, J.A., Smith, G.C., et al. (2009). Translational reprogramming following UVB irradiation is mediated by DNA-PKcs and allows selective recruitment to the polysomes of mRNAs encoding DNA repair enzymes. Genes Dev 23, 1207–1220. 10.1101/gad.516509.

50. Martinez, N.M., Su, A., Burns, M.C., Nussbacher, J.K., Schaening, C., Sathe, S., Yeo, G.W., and Gilbert, W.V. (2022). Pseudouridine synthases modify human pre-mRNA co-transcriptionally and affect pre-mRNA processing. Mol Cell 82, 645–659.e649. 10.1016/j.molcel.2021.12.023.

51. Hundley, H.A., and Bass, B.L. (2010). ADAR editing in double-stranded UTRs and other noncoding RNA sequences. Trends Biochem Sci 35, 377–383. 10.1016/j.tibs.2010.02.008.

52. Licht, K., Hartl, M., Amman, F., Anrather, D., Janisiw, M.P., and Jantsch, M.F. (2019). Inosine induces context-dependent recoding and translational stalling. Nucleic Acids Res 47, 3–14. 10.1093/nar/gky1163.

53. Nishikura, K. (2016). A-to-I editing of coding and non-coding RNAs by ADARs. Nat Rev Mol Cell Biol 17, 83–96. 10.1038/nrm.2015.4.

54. Ling, S.C., Polymenidou, M., and Cleveland, D.W. (2013). Converging mechanisms in ALS and FTD: disrupted RNA and protein homeostasis. Neuron 79, 416–438. 10.1016/j.neuron.2013.07.033.

55. Jonkers, I., and Lis, J.T. (2015). Getting up to speed with transcription elongation by RNA polymerase II. Nat Rev Mol Cell Biol 16, 167–177. 10.1038/nrm3953.

56. Shi, M., Zhang, H., Wang, L., Zhu, C., Sheng, K., Du, Y., Wang, K., Dias, A., Chen, S., Whitman, M., et al. (2015). Premature Termination Codons Are Recognized in the Nucleus in A Reading-Frame Dependent Manner. Cell Discov 1. 10.1038/celldisc.2015.1.

