## supplementary Figure for "Translation sustains productive Pol II elongation through maintenance of nuclear RNA surveillance"

### Supplement Figure 1

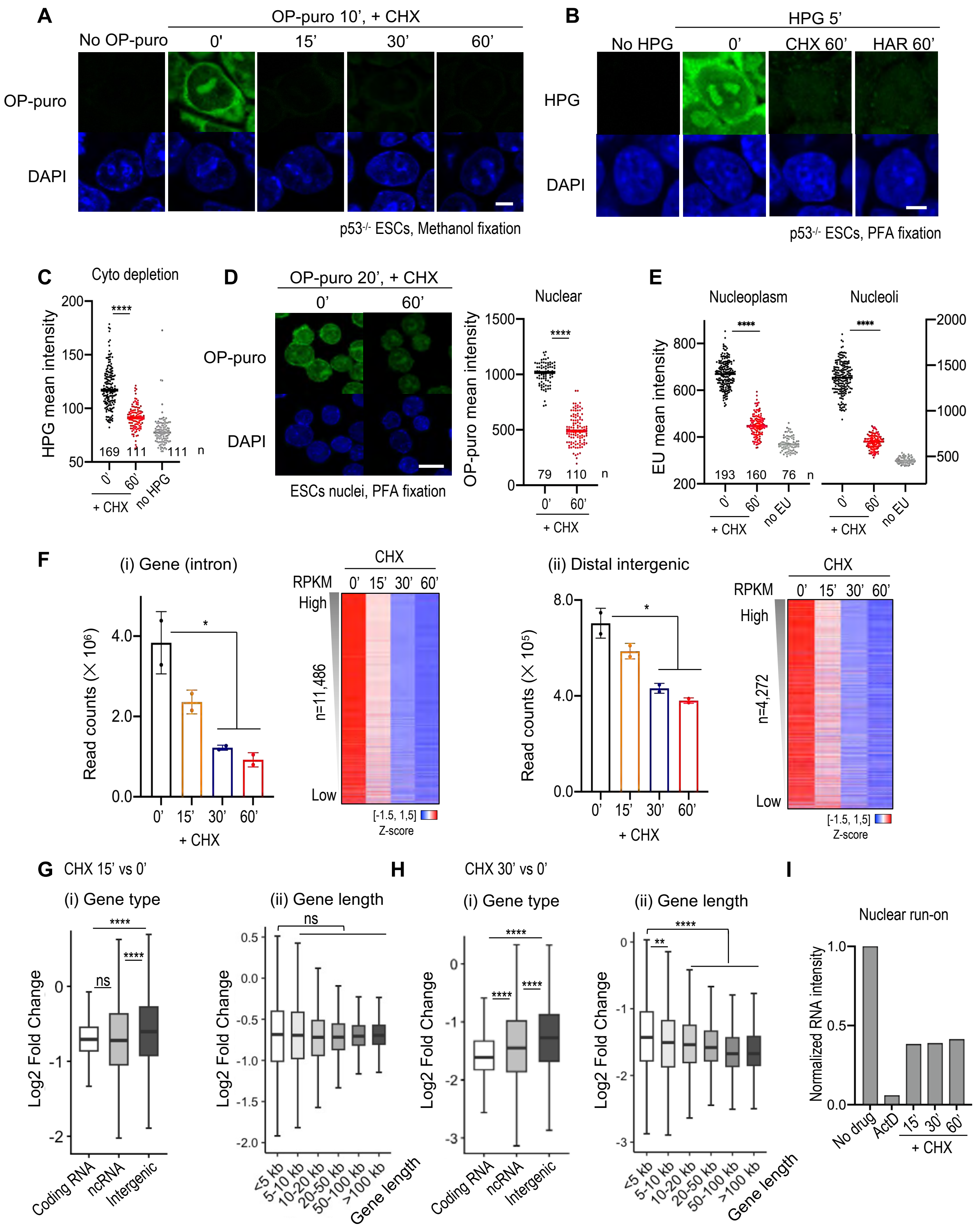

### Supplementary Figure 2

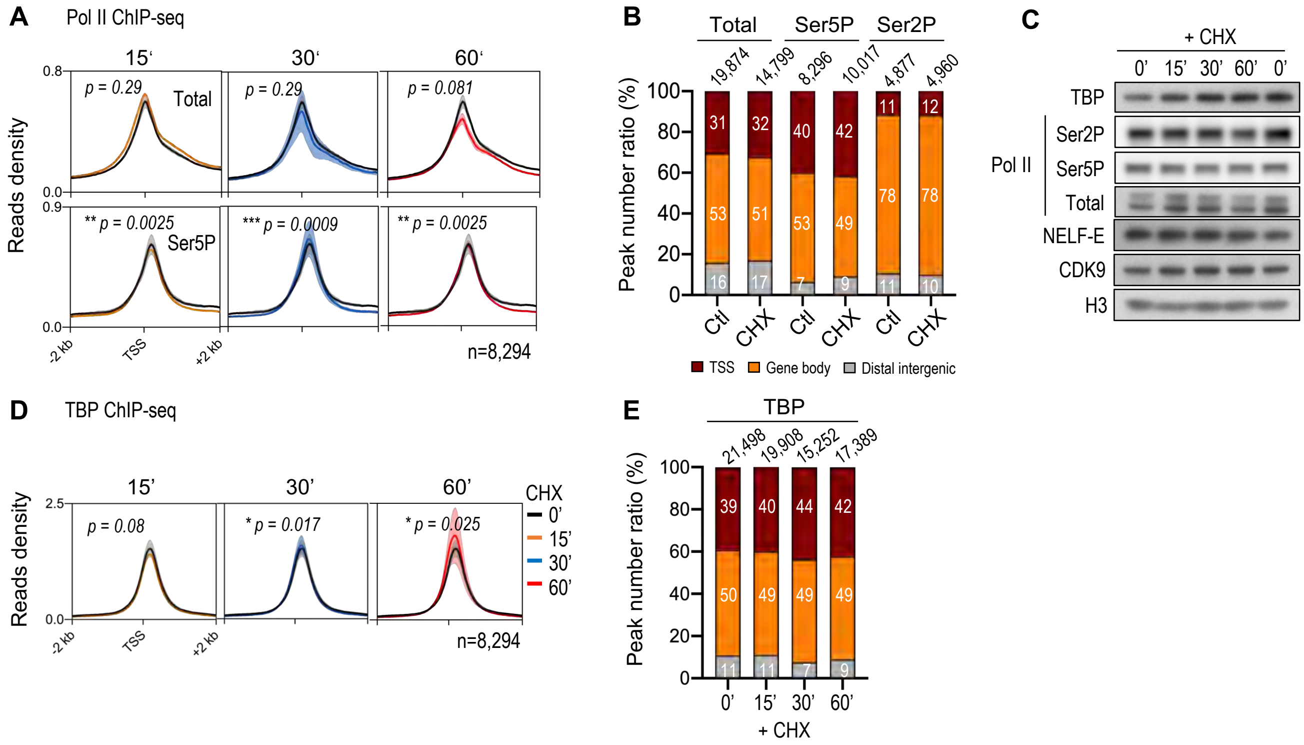

### Supplementary Figure 3

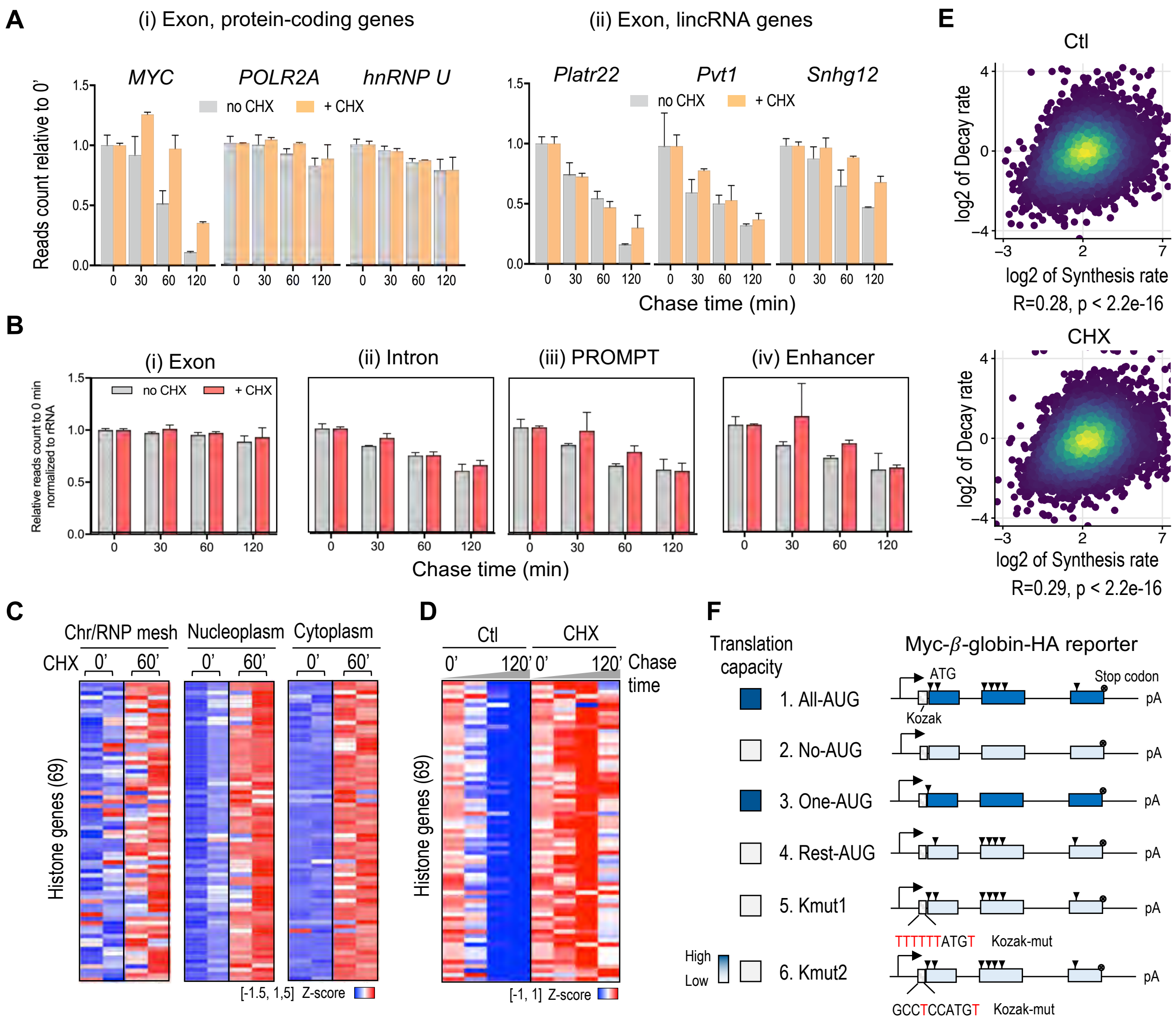

### Supplementary Figure 4

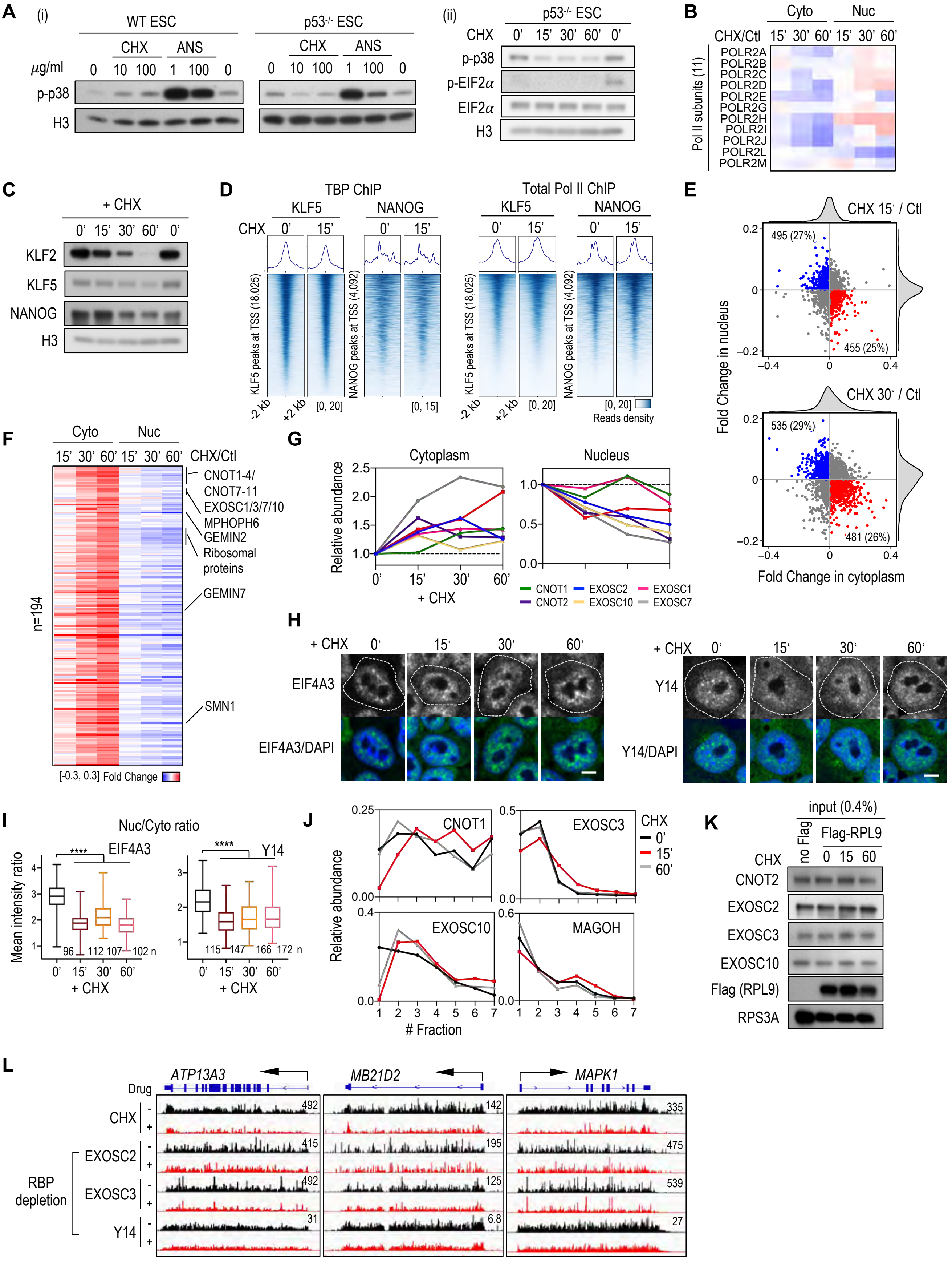

### Supplementary Figure 5

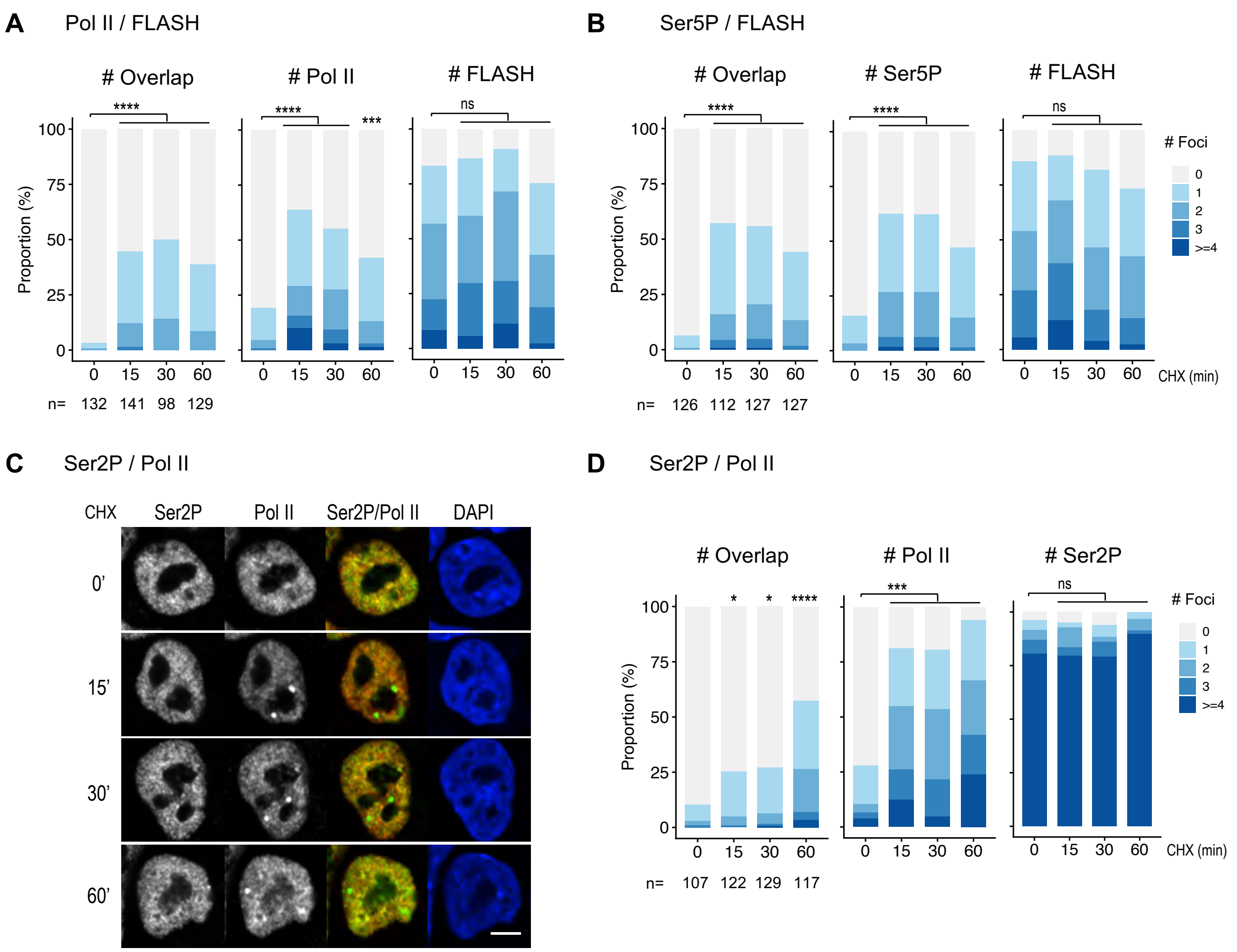

**Supplement Figure 1 Acute translation inhibition attenuates nascent transcription, related to Fig. 1**

(A) OP-puro incorporation for translational output. OP-puro were labeled for 10 min, followed by fixation, click chemistry, and fluorescence imaging. CHX treatment efficiently reduced OP-puro-labeled nascent peptides, indicating inhibition of translation. Scale bar: 5 $\mu$m.

(B) HPG incorporation for translational output. HPG were labeled for 5 min, followed by fixation, click chemistry, and fluorescence imaging. CHX and HAR treatment efficiently reduced HPG-labeled nascent peptides, indicating inhibition of translation. Scale bar: 5 $\mu$m.

(C) Quantification of nuclear HPG intensity in Fig. 1A. The number of nuclei analyzed is shown below. Statistical significance was assessed using a two-sided Wilcoxon rank-sum test.

(D) OP-puro incorporation for translational output in isolated nuclei. Isolated nuclei were incubated with OP-puro at 30 °C for 20 min, followed by fixation, click chemistry, and fluorescence imaging. CHX treatment reduced OP-puro-labeled nascent peptides in isolated nuclei. This suggests that OP-puro incorporation in isolated nuclei was reduced by CHX treatment. Quantification of nuclear OP-puro intensity is shown on the right. The number of nuclei analyzed is shown below. Statistical significance was assessed using a two-sided Wilcoxon rank-sum test. Scale bar: 10 $\mu$m.

(E) Quantification of EU intensity in nucleoli and nucleoplasm in Fig. 1B. The number of nuclei analyzed is shown below. Statistical significance was assessed using a two-sided Wilcoxon rank-sum test.

(F) Bar plot and heatmap showing EU-seq (3 min labeling) total read counts (left panel) and expression level of each bin (right panel) across intronic regions (i) and intergenic bins (ii) following 15-60 min CHX treatment. Intronic regions of each gene were used to quantify the nascent RNA. Distal intergenic regions were defined as genomic intervals located >10 kb away from annotated genes in the mm10 mouse genome. Compared with distal intergenic regions, which exhibit more stochastic transcription, genic regions decreased to ~60% of baseline levels within 15 min of CHX treatment, indicating higher sensitivity of productive transcription to translation inhibition. Bars represent two biological replicates and heatmap are shown as Z-score normalized values. Genes or bins with no detectable reads across all conditions were excluded.

(G-H) Box plots showing EU-seq signal changes (3 min labeling) following 15 or 30 min CHX treatment across genomic regions (coding RNA, ncRNA, and intergenic bins; n = 10,019/467/4,272) and gene length categories (<5, 5–10, 10–20, 20–50, 50–100, >100 kb). Gene numbers at each time point for each length category were: 15 min (n = 2523/1827/2838/4209/2294/2448) and 30 min (n = 2463/1803/2817/4179/2294/2451). Boxes indicate the $25^{th}$ - $75^{th}$ percentiles with medians shown, and whiskers represent the $5^{th}$ -$95^{th}$ percentiles. Outliers were omitted. Statistical significance was assessed using two-sided Wilcoxon rank-sum tests. Statistical comparisons were made between each group and the first group (<5 kb).

(I) Quantification of nuclear transcription signals normalized to *18S/28S* rRNA.

Significance: ns (>0.05), * (≤0.05), ** (≤0.01), *** (≤0.001), **** (≤0.0001). The same criteria were applied to all analyses shown in this study.

**Supplement Figure 2 Reduced Pol II elongation efficiency without impaired initiation, related to Fig. 2**

(A) Metagene analysis of Total, Ser5P and Ser2P Pol II ChIP-seq signals across 8,294 CHX-responsive genes (significantly changed during 15-60 min CHX treatment). Profiles are shown at genes transcriptional start sites (TSSs) with ±2 kb flanking regions. Solid lines indicate mean values of two biological replicates and shaded areas represent SEM. Statistical significance was assessed using a two-sided Kolmogorov-Smirnov test. Signals were normalized by sequencing depth.

(B) Bar plot showing the genome-wide peak distribution of Total, Ser5P, and Ser2P Pol II ChIP-seq signals. The total number of peaks is shown above each bar. No substantial difference in genomic Pol II peak distribution was observed between control and CHX-treated groups.

(C) Western blot analysis of total cellular protein levels following CHX treatment.

(D) Metagene analysis of TBP ChIP-seq signals across 8,294 CHX-responsive genes. Profiles are shown at genes TSSs with ±2 kb flanking regions. Solid lines indicate mean values of two biological replicates and shaded areas represent SEM. Statistical significance was assessed using a two-sided Kolmogorov-Smirnov test. Signals were normalized by sequencing depth.

(E) Bar plot showing the genome-wide peak distribution of TBP Pol II ChIP-seq signals. The total number of peaks is shown above each bar. No substantial difference in genomic peak distribution was observed between control and CHX-treated groups.

**Supplement Figure 3 Attenuation of nuclear RNA decay concurrent with reduced RNA synthesis, related to Fig. 3**

(A) Bar plots showing exonic read counts for representative genes. MYC and lincRNA transcripts showed lower baseline stability, while POLR2A and hnRNP U transcripts were relatively stable during the 120-min chase. CHX treatment broadly stabilized all measured transcripts. Values were normalized to rRNA and then to the 0 min chase time for visualization.

(B) Bar plot showing sum read counts across exonic, intronic, PROMPT, and enhancer regions. Bar plot showing summed read counts across exonic, intronic, PROMPT, and enhancer regions. CHX treatment led to global stabilization of all transcript classes.

(C) Heatmap showing steady-state RNA levels of 69 replication-dependent histone genes across chromatin/RNP mesh (Chr/RNP mesh), nucleoplasm, and cytoplasm fractions. Values represent two biological replicates and are Z-score normalized. Data related to Fig. 3C.

(D) Heatmap showing RNA stability of 69 replication-dependent histone genes measured by EU pulse-chase experiments. Gene body read counts of histone genes were normalized to rRNA, expressed relative to the 0 min time point, and Z-score transformed for visualization. Data related to Fig. 3F.

(E) Dot plot showing the correlation between RNA synthesis and nuclear degradation rate with or without CHX treatment. Pearson correlation and t-test were used for statistical analysis. Data related to Fig. 3G.

(F) Schematic of the β-globin reporter system according to previous study[56]. A series of ORF variants were constructed to systematically modulate translational potential by altering start codon and initiation context, including constructs with all AUGs intact (All-AUG, same as wild-type sequence), all in-frame AUG start codons mutated (No-AUG), or only the first AUG retained with all downstream AUGs mutated (One-AUG), selective mutation of the first AUG (Rest-AUG), and disruption of Kozak consensus sequences (Kmut1 and Kmut2). Reporter constructs were cloned into expression vectors and stably introduced into cells to generate polyclonal reporter cell lines. Related to Fig. 3H.

**Supplement Figure 4 Nuclear RNA surveillance protein redistribution in the absence of ribotoxic stress or pluripotency loss, related to Fig. 4**

(A) Western blot analysis of p38 and EIF2α in total cellular lysates. Wild-type and $p53^{-/-}$ ESCs were treated with CHX or HAR at the indicated concentrations and times.

(B) Heatmap showing fold changes in Pol II subunit abundance relative to control, based on quantitative mass spectrometry.

(C) Western blot analysis of pluripotency transcription factors in total cellular lysates. $p53^{-/-}$ ESCs were treated with CHX for the indicated times.

(D) Heatmap showing TBP and Total Pol II ChIP distribution across KLF5- and NANOG-target genes and their ±2 kb flanking regions. Signals represent the mean of two biological replicates and are Z-score normalized. Genes or bins with no detectable reads across all conditions were excluded.

(E) Dot plot showing nucleocytoplasmic redistribution of proteins following 15–30 min CHX treatment, based on quantitative mass spectrometry. Proteins with increased cytoplasmic and decreased nuclear abundance are shown in red, whereas those with the opposite pattern are shown in blue. Related to Fig. 4A.

(F) Heatmap showing fold changes in protein abundance relative to control. Highlighted proteins increased by >10% in the cytoplasm at 60 min CHX treatment and showed a corresponding decrease in nuclear abundance (n = 194 of 452). Some of RNA catabolic and snRNP assembly factors are indicated on the right. Data related to Fig. 4A.

(G) Quantification of band intensities of representative proteins in cytoplasmic and nuclear fractions relative to 0 min. Related to Fig. 4D.

(H) Immunofluorescence analysis of EIF4A3 and Y14 distribution following CHX treatment for 15-60 min. EIF4A3 was detected using anti-Flag antibodies in ESCs expressing endogenous Flag–EIF4A3, and Y14 was detected using a specific antibody. Cell boundaries are outlined with dashed gray lines. EIF4A3 and Y14 were retained in the cytoplasm after CHX treatment. Scale bars, 5 $\mu$m.

(I) Quantification of immunofluorescence intensity in Supplementary Fig. 4H. Nuclear-to-cytoplasmic intensity ratios were calculated to quantify protein redistribution. The number of cells analyzed is shown below. Statistical significance was assessed using a two-sided Wilcoxon rank-sum test for each treatment time point versus control.

(J) Quantification of band intensities of representative proteins from polysome profiling. Relative abundance represents the ratio of protein signal in each fraction to total signal across seven fractions. Related to Fig. 4F.

(K) Western blot analysis of the input fraction from RPL9 co-immunoprecipitation experiments. Related to Fig. 4G.

(L) Representative IGV tracks of EU-seq signals at the *ATP13A3*, *MB21D2*, and *MAPK1* loci under CHX treatment or upon depletion of EXOSC2, EXOSC3, and Y14. EU labeling was performed for 3 min in CHX-treated samples and for 10 min in EXOSC2-, EXOSC3-, or Y14-depleted conditions. Tracks represent merged biological replicates.

**Supplement Figure 5 Accumulation of Pol II initiation foci at histone locus bodies, related to Fig. 5**

(A) Quantification of Pol II and FLASH foci and their overlap. The number of cells analyzed is shown below. Statistical significance was assessed using pairwise Fisher's exact tests with FDR correction. Statistical comparisons shown are between CHX-treated and control groups.

(B) Quantification of Ser5P Pol II and FLASH foci and their overlap. The number of cells analyzed is shown below. Statistical significance was assessed using pairwise Fisher's exact tests with FDR correction. Statistical comparisons shown are between CHX-treated and control groups.

(C) Immunofluorescence showing CHX-induced redistribution of Ser2P and Total Pol II. Scale bar: 5 $\mu$m.

(D) Quantification of Ser2P and Total Pol II foci and their overlap. The number of cells analyzed is shown below. Statistical significance was assessed using pairwise Fisher's exact tests with FDR correction. Statistical comparisons shown are between CHX-treated and control groups.
